# Loss of clustered protocadherin diversity alters the spatial distribution of cortical interneurons

**DOI:** 10.1101/2020.10.08.331488

**Authors:** Nicholas Gallerani, Edmund Au

**Affiliations:** Department of Pathology & Cell Biology, Columbia University Irving Medical Center, New York NY, 10032; Department of Rehabilitative Medicine and Regeneration, Columbia University Irving Medical Center, New York NY, 10032; Columbia Translational Neuroscience Initiative Fellow

## Abstract

Cortical interneurons (cINs) are locally-projecting inhibitory neurons that are distributed throughout the cortex. Due to their relatively limited range of influence, their arrangement in the cortex is critical to their function. cINs achieve this arrangement through a process of tangential and radial migration, and apoptosis during development. In this study, we investigated the role of clustered protocadherins (cPcdhs) in establishing the spatial patterning of cINs. cPcdhs are expressed in cINs, and are known to play key functions in cell spacing and cell survival, but their role in cINs is poorly understood. Using spatial statistical analysis, we found that the two main subclasses of cINs, parvalbumin-expressing (PV) and somatostatin-expressing (SST) cINs, are non-randomly spaced within subclass, but randomly with respect to each other. We also found that the relative laminar distribution of each subclass was distinctly altered in whole α- or β-cluster mutants. Examination of perinatal timepoints revealed that the mutant phenotypes emerged relatively late, suggesting that cPcdhs may be acting during cIN morphological elaboration and synaptogenesis. We then analyzed an isoform-specific knockout for pcdh-αc2 and found that it recapitulated the α-cluster knockout, but only in SST cells, suggesting that subtype-specific expression of cPcdh isoforms may help govern subtype-specific spatial distribution.

## Introduction

The correct spatial distribution of neurons can be critical to brain function. This is true in the case of cortical interneurons (cINs) of the neocortex, which are locally projecting inhibitory cells that carry out key processes, including controlling cortical rhythmicity (Sohal VS et al. 2009), boosting signal salience (Lee SH et al. 2012; Song YH et al. 2020), and regulating spike timing (Tiesinga P et al. 2008). Diverse subclasses of cINs are found throughout the neocortex, where they establish repetitive microcircuit motifs (Tremblay R et al. 2016).

How interneuron subclasses appropriately distribute and establish an inhibitory network poses a difficult logistical problem. Developmentally, cIN arise from distal progenitor domains and migrate long distances into the cortex (Wamsley B and G Fishell 2017). Following migration, cIN number is reduced through apoptotic cell death (Southwell DG et al. 2012) and cINs extend complex, branched dendritic and axonal arbors that overlap considerably with neighboring cINs, enabling a single cIN to inhibit multiple pyramidal cells (PC), and for a single PC to be innervated by multiple cINs (Fino E and R Yuste 2011; Packer AM et al. 2013). How these stages are orchestrated to allow cINs to establish a distributed network of inhibition is poorly understood.

An appealing molecular candidate that could mediate this process is a family of adhesion molecules known as clustered protocadherins (cPcdhs). cPcdhs are arranged in 3 gene clusters that encode 58 distinct isoforms of alpha-(α), beta-(β) and gamma-(γ) cPcdhs (Chen WV and T Maniatis 2013). They impart neurons with single cell identity by differential (and largely stochastic) expression of cPcdhs, which form combinatorial cPcdh recognition complexes (Esumi S et al. 2005; Hirano K et al. 2012; Brasch J et al. 2019). These molecules mediate crucial circuit formation functions (Mountoufaris G et al. 2017) including dendritic self-avoidance (Lefebvre JL et al. 2012), axonal tiling (Chen WV et al. 2017), and cell survival (Hasegawa S et al. 2016; Garrett AM et al. 2019). Deficits in α-cPcdh diversity can cause detectable changes in sensory processing (Yamagishi T et al. 2018). cPcdhs are expressed in cINs (Hirano K *et al*. 2012; Mancia Leon WR et al. 2020), and recently, it was demonstrated that γ-cPcdhs regulate programmed cell death in cINs (Mancia Leon WR *et al*. 2020). However, the role of α- and β-cPcdhs in cINs is still poorly understood. We therefore sought to determine if α- and β-cPcdhs are important for other aspects of cIN development, such as facilitating proper subclasses spatial distribution.

To test this, we developed a method of registering cIN cell body positions in XY coordinate space. This allowed us to analyze spatial distribution of the two main interneuron subclasses, parvalbumin+ (PV) and somatostatin+ (SST) cINs. Together, they account for ∼70% of all cINs (Tremblay R *et al*. 2016) and share a developmental origin in the medial ganglionic eminence (MGE) (Wamsley B and G Fishell 2017). The density of cIN processes makes them difficult to directly assess at the population level *in vivo*, but we reasoned that since interneurons possess locally-projecting processes, their cell body placement was a reasonable and accessible proxy for the area of influence of the neuron. The position of the cell body has been similarly used to study neuron spacing in locally projecting inhibitory retinal amacrine cells (Fuerst PG et al. 2008).

Using this method, we compared cINs in wildtype (WT) mice, whole α-cPcdh knock-out (α-KO) and whole β-cPcdh knock-out (β-KO) mutant mice and found that loss of cPcdh diversity differentially affects PV and SST cINs. In the adult primary visual cortex (V1) of the α-KO, we found that PV cell density was reduced, while SST density was not. Additionally, we observed subclass-specific alterations in laminar distribution of PV and SST populations. Both α- and β-KOs exhibited defects, but the phenotype was most pronounced in V1 of the α-KO. Focusing on V1 in WT and α-KO, we performed an analysis at perinatal timepoints and found that the mutant phenotype arises after migration and during the late phase of canonical cell death. To further understand how the α-KO differentially affects cIN, we examined pcdh-αc2 (αc2), an α-cPcdh isoform which is non-stochastically expressed (Chen WV and T Maniatis 2013), and expressed at high levels in both PV and SST in adult V1 (Supp8). In the αc2-specific mutant (αc2-KO), the SST phenotype was similar to the α-KO phenotype, but PV cells were unaffected. Taken together, our work suggests that cPcdhs play an important role in the laminar distribution of cINs during perinatal development. Further, our work suggests that subclasses of cINs employ specific cPcdh isoforms (αc2 in the case of SST cells) as they establish a distributed inhibitory network in the cortex.

## Materials and Methods

### Animals

All animal experiments were performed in compliance with established protocols approved by IACUC. α-KO and β-KO mice were generated according to methods described previously (Mountoufaris G *et al*. 2017). For the α-KO mouse, all variable exons in the α-cluster were deleted while the constant exons were left intact. For the β-KO mouse, the entire β-cluster was deleted. The αc2-KO was generated according to methods described in a previous study (Chen WV *et al*. 2017), but only ablates the Pcdh-αc2 variable exon, rather than both Pcdh-αc1 and Pcdh-αc2. For perinatal experiments, α-KO mice were crossed with BAC-Nkx2.1-Cre; Ai9 mice (<u>Jackson</u>) (<u>Jackson</u>). We analyzed the following number of mice: (WT for whole cluster: n=7, α-KO: n=10, β-KO: n=8, WT for αc2-KO: n=5, αc2-KO: n=5, P7 Nkx2.1Cre;Ai9;α-WT: n=6, P7 Nkx2.1Cre;Ai9; α-KO: n=6, P13 Nkx2.1Cre;Ai9; α-WT: n=5, Nkx2.1Cre;Ai9; α-KO: n=5). All mice were maintained in a mixed 129 and C57BL/6J genetic background.

### Histology

Mice were anesthetized with ketamine (87.5 mg/kg) xylazine (12.5mg/kg) via intraperitoneal injection and transcardially perfused with phosphate buffered saline (PBS) followed by 4% paraformaldehyde (PFA) in PBS. Brains were post-fixed in 4% PFA/PBS overnight at 4°C, embedded in low melt agarose and then parasagitally sectioned on a Leica VT1000S vibratome (50µm thickness). Sections were stored at −30°C in an antifreeze solution containing polyethylene glycol, glycerol, and PBS.

### Immunohistochemistry

Sections were transferred from antifreeze solution to PBS and washed 3x in PBS. Heat-induced epitope retrieval was then performed by incubating sections in 10mM sodium citrate (pH 8.5) at 80°C for 30 minutes. Sections were then incubated at room temperature in a blocking/permeabilization buffer consisting of 2% w/v nonfat-dried milk, 0.3% TritonX, and 0.01% sodium azide in PBS. Sections were incubated in primary antibodies diluted in 2% normal donkey serum and 0.02% Tween20 in PBS for 24-72 hours at 4°C. Primary antibodies used in this study: guinea pig anti-parvalbumin (1:750, Immunostar), rat anti-somatostatin (1:300, EMD Millipore), rabbit anti-active caspase 3 (1:1000, Sigma-Aldrich), rabbit anti-serotonin transporter (1:1000, Immunostar). Sections were incubated in secondary antibodies diluted 1:1000 in PBS for 2 hours at room temperature. All secondary antibodies were Alexa Fluor conjugated affinity-purified IgG raised in donkey host (Jackson). Sections were incubated in 300nM 4’,6-diamidino-2-phenylindole (DAPI) diluted in PBS for 15 minutes at room temperature. Sections were mounted on SuperFrost Plus slides (<u>Fisher Scientific</u>) using Fluoromount-G (<u>Southern Biotech</u>) and #1.5 cover slips.

### Imaging

Adult brain tissue was imaged on a CSU-W1 spinning disk confocal microscope (<u>Yokogawa</u>). 20-30µm Z-stacks were acquired at 2µm Z-intervals using 10x or 20x objectives. Image fields were tiled and stitched in NIS-Elements (<u>Nikon</u>). Median intensity projection images were generated (<u>ImageJ</u>) from 3-4 consecutive optical slices to minimize unwanted signal from cells located deeper in the tissue outside of the focal plane. Tiled images of sagittal sections were assigned an “Allen Brain Atlas Number” from 1 to 21 based on the P56 Sagittal Reference Atlas (<u>Allen Institute</u>). Based on atlas number and stereotaxic coordinates (Paxinos and Franklin, *The mouse brain in stereotaxic coordinates*) images containing specific functional regions of interest (ROI) such as V1 and somatosensory cortex were identified for analysis. Perinatal brain tissue was imaged on a <u>Zeiss Axio Imager M2</u> epifluorescence microscope. In all cases, at least three images of comparable ROIs were captured per mouse.

### Semi-Automated Cell Quantification

Raw images were processed for quantification using a series of custom <u>ImageJ macro scripts</u>. To blind the experimenter, images were assigned a random code using an existing ImageJ macro script (<u>Filename Randomizer, Tiago Ferreira, 2009</u>). Multichannel images were split into single channels. DAPI was used to manually specify layers drawn onto the image using ImageJ selection tools, on an ROI roughly 1mm in width. L1 was not included in the analysis because PV and SST interneurons are absent in this layer. Cortical layers were visually distinguishable from each other based on differences in density of DAPI stained nuclei. Line selections were manually drawn onto the image to demarcate the boundaries between each layer. The XY coordinates of the area and line selections were stored.

Single channel images containing cells were processed with a custom ImageJ pipeline incorporating standard image filtering and automatic thresholding tools built into ImageJ. A difference of Gaussians (DoG) filter was applied to the image. This filter applies a Gaussian blur to the original image, with σ_1_ slightly smaller than the smallest cells in the image. Then, a second Gaussian blur with σ_2_=2*σ_1_ was applied to the original image. The resulting blurred image is subtracted from the first less blurred image, and results in an image where desired large bright objects, such as cells, are enhanced, while undesired small bright objects, such as fine staining of PV positive neuronal projections, are suppressed. In order to correct brightness variability across the image, we applied contrast limited adaptive histogram equalization (CLAHE). Finally, we applied the default automatic thresholding algorithm built into ImageJ to obtain a binarized version of the original image. Then, we processed the binarized image with a watershed algorithm and selected particles based on size and circularity to obtain a final set of automated cell counts. A final manual quality control step was performed to ensure cells were counted accurately. After this step, the XY coordinates of PV and SST cells were stored.

### Spatial Statistics

Once all XY coordinates of regions, layers, and cells were obtained, spatial measurements such as cortical thickness, cell density, nearest neighbor distance (NND), variance of the NND, and paired correlation function were computed using <u>Spatstat (Baddeley AT, R. 2005)</u>, an open-source platform for analyzing spatial point patterns in R.

#### Cortical Dimensions

Cortical thickness was calculated for each image by measuring lines spaced at 1µm intervals along the axis perpendicular to the bottom of layer 6 and calculating the median of the distribution of lines. We generated points at 1µm intervals along the bottom of layer 6 and then generated the same number of points spread equally along the top of layer 2/3, and calculated the pairwise distance between each pair of points. This same method was used to calculate the thickness of each layer.

#### Cell Density

We calculated cell density by dividing the number of cells counted by the measured area. The fraction of cells in a given layer was determined by calculating the number of cells that were between the layer annotation lines and dividing by the total number of cells in the ROI.

#### Nearest Neighbor Distance

The NND is the distance between a given cell and its closest cell. This calculation was performed between the same cell types as well as different cell types. NND was also used to determine double positive cells. Cells of different types that had NNDs smaller than the measured radius of the cell body were considered double positive.

The variance of the NND can be used to measure the regularity of spacing between cells. Cells with lower variance are more regularly spaced, while a higher variance indicates that they are more randomly spaced. In order to account for differences in mean NND which may affect the variance, we divided the standard deviation of the NND by the mean NND to give a dimensionless ratio known as the coefficient of variance (CV). This allowed us to directly compare the regularity in spacing between groups which had different mean NND.

#### Paired Correlation Function

To measure the degree of spatial clustering or inhibition we used the pair correlation function g(*r*), which is the ratio of observed cells at a given distance *r* from a reference cell to the expected cells at distance *r* if the distribution were random. In a completely random distribution, the density of cells at any *r* will be equal to the density of the entire field, so g(*r*)=1. Values below 1 indicate spatial inhibition, while values above 1 indicate spatial clustering. g(*r*) for all cells in an ROI is calculated using kernel smoothing. g(*r*) for each mouse was calculated by averaging each ROI. g(*r*) values for genotypes were the average of the mice.

### Statistical Significance

Statistical significance values were calculated in <u>GraphPad Prism 8</u>. Statistical tests and p values are listed in figure legends. Significance values are reported as following: *=p<0.05, **=p<0.01, ***=p<0.001. Post-hoc power analysis was performed using <u>G*Power 3.1.9.7</u>

## Results

### Parvalbumin (PV) and somatostatin (SST) expressing interneurons are spatially independent populations

To characterize the spatial arrangement of cINs in normal and mutant animals, we first established a pipeline to convert raw image data into a form that could be readily analyzed using the spatial analysis software suite, Spatstat (Baddeley AT, R. 2005) (Fig1A-B, Supp1). This method allowed us to analyze a number of different spatial characteristics of cIN classes. For our analysis, we focused on two cortical regions in which cINs have been well-studied: primary visual (V1) and primary somatosensory (S1) cortices.

**Figure 1.**
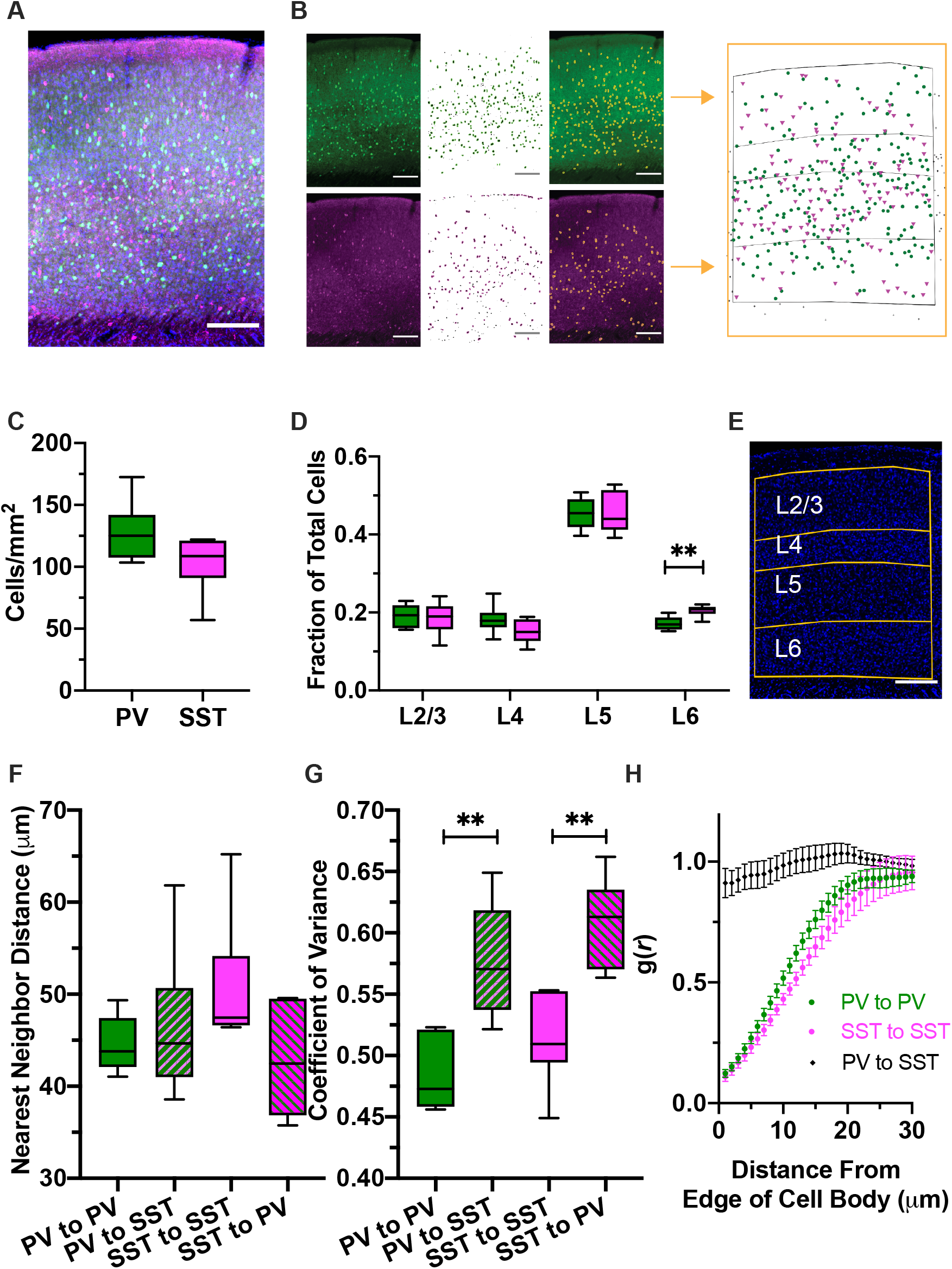
Parvalbumin (PV) and somatostatin (SST) expressing cortical interneurons (cINs) are spatially independent populations. A-Representative fluorescent image of P30 WT primary visual cortex (V1) labeled with DAPI (blue), anti-PV(green), and anti-SST (magenta), with cortical layers obtained from DAPI image (yellow); scale bar=200μm B-Raw images of PV (left top) and SST (left bottom) are processed to obtain XY coordinates of the cells (far right). See Supp 1 for more detail. C-Cell density (cells/mm^2^) of PV and SST neurons. PV neurons tend to be denser than SST neurons (p=0.0629, unpaired T-test with Welch’s correction) D-Relative proportion of PV and SST neurons in each cortical layer in V1, WT. PV and SST neurons are distributed similarly, but there are relatively more SST neurons in L6 (p=0.0095, 2-way ANOVA with Sidak’s multiple comparisons test (SMCT)) E-Representative image of DAPI (blue) stained tissue with layers of the cortex marked (yellow) F-Nearest neighbor distance (NND) between PV-PV, PV-SST, SST-SST, and SST-PV pairs in V1, WT. NND was not significantly different for PV-PV, PV-SST, SST-SST, or SST-PV pairs (p=0.111, 1-way ANOVA) G-Coefficient of variance (CV) of the NND for PV-PV, PV-SST, SST-SST, and SST-PV pairs in V1, WT. CV was significantly lower between PV-PV pairs compared to PV-SST pairs (p=0.0049, 1-way ANOVA with Dunnett’s (D) MCT), and SST-SST pairs compared to SST-PV pairs (p=0.0026) H-Paired correlation function (PCF) of PV-PV pairs (green), SST-SST pairs (magenta), and PV-SST pairs (black). Distance between PV-SST pairs is random, but distance between like-cell types is statistically non-random at relatively small distances (PV-PV: p<0.05 up to 19μm from edge of cell body, SST-SST: p<0.05 up to 19μm; 2-way ANOVA with SMCT).

Consistent with existing literature (Rudy B et al. 2011), PV neurons were about 20% more dense than SST neurons in V1 (Fig1C). Both types of neuron were found in every layer besides L1, but the majority of both types are found in L5 (Fig1D). Both types of neurons have largely similar laminar distribution patterns but SST neurons are relatively more abundant in L6 compared to L5 (Fig1D). S1 had no significant differences compared to V1 for all measurements other than cortical thickness, which was generally thicker in S1 (Supp2).

PV and SST cells share a common developmental origin (Wamsley B and G Fishell 2017), and largely occupy the same cortical space. We next examined how PV and SST cells are spaced within and between subclasses. The nearest neighbor distance (NND) between PV-PV pairs was not different than the NND between PV-SST pairs (Fig1F). This was also true for SST-SST and SST-PV pairs. To compare the regularity in spacing between groups, we used the coefficient of variance (CV) of the NND, which is a normalized measure of variance. A lower CV suggests greater regularity of spacing. The CV was significantly lower in PV to PV pairs compared to PV to SST pairs, as well as for SST to SST pairs compared to SST to PV pairs (Fig1G). This suggests greater regularity (even spacing) between groups compared to across groups. To see if cells exhibited spatial inhibition or clustering, we used the pair correlation function (PCF), g(*r*) (see Methods-Spatial Statistics-Paired Correlation Function). We found that matching cell type pairs had significantly smaller g(*r*) values than expected at modest distances from their cell bodies, while unmatched pairs were not significantly different than random at any distance (Fig1H). These data suggest that PV cells and SST cells distribute spatially independent of each other, but maintain spacing relative to homotypic cells.

### Loss of α-protocadherins reduces the density of PV neurons and alters the laminar distribution of both PV and SST neurons

To study how cPcdhs affect the spatial arrangement of cINs we performed the same measurements in the context of cPcdh mutants. We studied mice in which either all variable exons of the α cluster are deleted (α-KO) or all exons of the β cluster are deleted (β-KO). We did not examine γ-cluster mutants (γ-KO) because they are perinatal lethal (Hasegawa S *et al*. 2016; Garrett AM *et al*. 2019; Mancia Leon WR *et al*. 2020), before mature cIN spatial distribution is established. Since loss of a cPcdh cluster would reduce the potential pool of cPcdhs any given cell could draw from for self- and cell-cell recognition, we hypothesized that it would result in disruptions to PV and SST cell distribution.

In V1 of the α-KO we found that the density of PV cells was significantly lower compared to WT (Fig2B). In every layer except L6, PV density was significantly reduced (Fig2F) while the density of SST was unaffected (Fig2B, G). The NND between PV pairs was also significantly increased in the α-KO (Fig2C), consistent with a reduction in cell density. Indeed, we found that the NND was significantly correlated with cell density for all cell types and genotypes analyzed (Supp3). We also noted a change in the relative distribution of PV cells, with relatively fewer PV cells in L5 and more PV cells in L6. (Fig2D). With SST cells (Fig2B, G), we noted changes in their relative laminar distribution: more of the SST cells were found in L2/3, and fewer in L5 (Fig3E). In S1, we also found an increase in the relative amount of L6 PV cells, but other measures were unaffected (Supp4). However, we did note trends that resembled the effects seen in L2/3 and L5 of V1 for SST cells, which were just outside the significance threshold (Supp4E). Altered laminar distribution in both PV and SST were not accompanied by changes in CV or PCF (Supp5).

**Figure 2.**
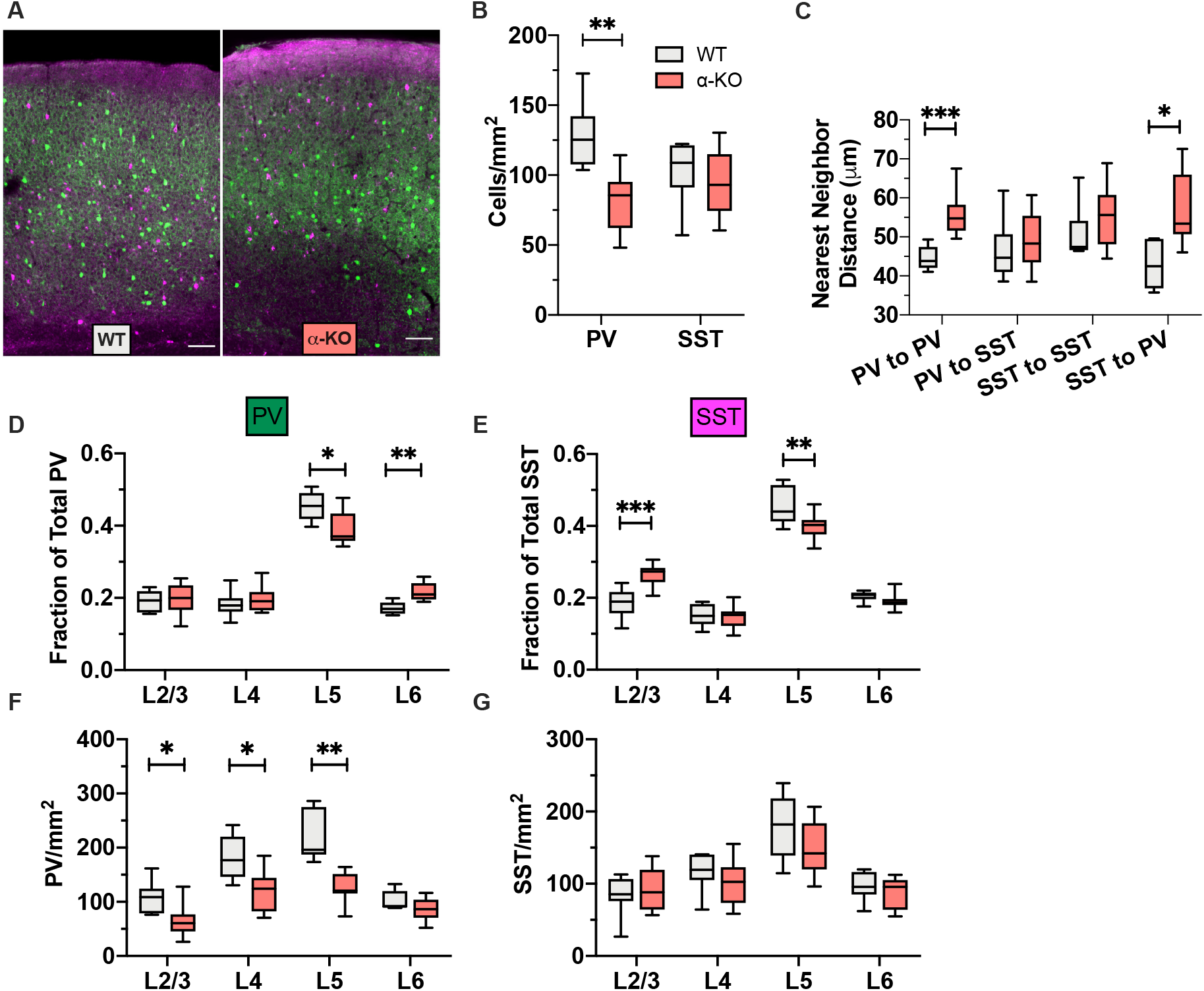
Loss of α-protocadherins reduces the density of PV neurons and alters the laminar distribution of both PV and SST neurons. A-Representative images of WT (left) and α-KO (right) V1, PV (green) and SST (magenta) cIN; scale bar=200μm B-PV and SST cell density in cells/mm^2^ for WT and α-KO V1 cortex. PV density is reduced while SST density is unchanged (PV: p=0.0033, SST: p=0.7022, 1-way ANOVA with DMCT) C-Nearest neighbor distance (NND) between PV-PV, PV-SST, SST-SST, and SST-PV pairs in WT and α-KO V1. NND between PV-PV pairs is increased. The distance between SST-PV pairs is also increased (Left to Right (LtR): p=0.0002, p=0.9801, p=0.7653, p=0.0134, 2-way ANOVA with SMCT). D-Relative proportions of PV cells occupying each cortical layer in V1, WT v α-KO. The relative amount of PV cells is reduced in L5 and increased in L6 (LtR: p=0.9176, p=0.6694, p=0.0146, p=0.0014, 2-way ANOVA with SMCT) E-Relative proportions of SST cells occupying each cortical layer in V1, WT v α-KO. The relative amount of SST cells is reduced in L5 and increased in L2/3 (LtR: p=0.0004, p=0.9797, p=0.0084, p=0.7596, 2-way ANOVA with SMCT) F-Density of PV cells in each cortical layer in V1, WT v α-KO. The density of PV cells is reduced in all layers except L6 (LtR: p=0.0309, p=0.0218, p=0.0027, p=0.2478, 2-way ANOVA with SMCT) G-Density of SST cells in each cortical layer in V1, WT v α-KO. The density of SST cells is unchanged in any layer (LtR: p=0.8043, p=0.5522, p=0.4019, p=0.6955, 2-way ANOVA with SMCT)

Differences in relative layer thickness between WT and a-KO were a potential confound. To assess this, we quantified cortical thickness, as well as the thickness of its layers. The cortex of the α-KO was significantly thicker overall and in L2/3 and L6 (Supp6A, B). However, layers were proportionally similar to WT (Supp6C), indicating that changes associated with L5 or L6 were not the result of the layers themselves being altered. We did find that L2/3 was modestly, but significantly greater (2%) as a relative portion of the α-KO cortex (Supp6C). In contrast, the difference in relative proportion of SST cells in L2/3 (19% in WT vs. 27% of SST in α-KO) was robust, suggesting that the change in relative cortical thickness in α-KO would be at most, a minor contributing factor.

### Loss of β-protocadherins alters the laminar distribution of PV and SST interneurons

Like the α-KO, we observed a significant reduction in the relative amount of PV cells in L5 (Fig3D), and also an upward trend in L6 (p=0.1050). PV laminar density was mostly unchanged (Fig3B, F). The mean density of SST cells was unchanged overall or in any layer (Fig3B, G). However, we observed a significant increase in the relative amount of SST cells in L2/3 (Fig3E), and a trend (p=0.1601) towards reduced SST in L5. Together, our analysis of the β-KO revealed a similar, but milder, phenotype compared with α-KO.

**Figure 3.**
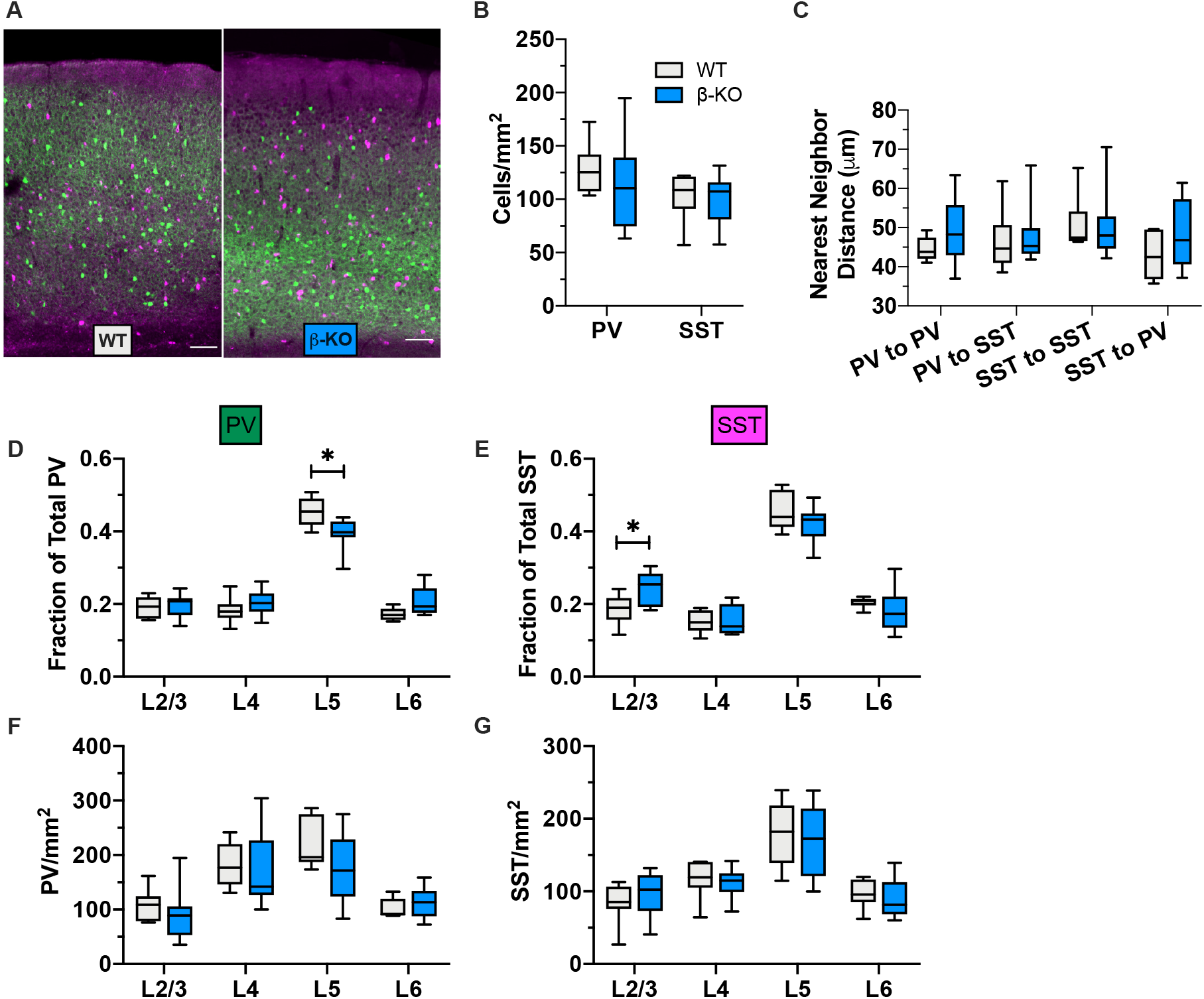
Loss of β-protocadherins alters the laminar distribution of PV and SST cIN. A-Representative images of WT (left) and β-KO (right) V1 PV (green) and SST (magenta) cIN; scale bar=200μm B-PV and SST cell density in cells/mm^2^ for WT and β-KO V1 cortex. PV and SST density is not different between WT and β-KO (PV: p=0.7932, SST: p=0.9882, 1-way ANOVA with DMCT) C-NND between PV-PV, PV-SST, SST-SST, and SST-PV pairs in WT and β-KO V1. NND was not altered for any group (LtR: p=0.5313, p=0.9996, p=0.9991, p=0.6664, 2-way ANOVA with SMCT) D-Relative proportions of PV cells occupying each cortical layer in V1, WT v β-KO. The relative amount of L5 PV cells was reduced (LtR: p=0.9655, p=0.5049, p=0.0326, p=0.1050, 2-way ANOVA with SMCT) E- Relative proportions of SST cells occupying each cortical layer in V1, WT v β-KO. The relative amount of L2/3 SST cells was increased (LtR: p=0.0164, p=0.9943, p=0.1601, p=0.5354, 2-way ANOVA with SMCT) F- Density of PV cells in each cortical layer in V1, WT v β-KO. The laminar density of PV cells was not changed compared to WT, but trended similar to the α-KO in L5 (LtR: p=0.7352, p=0.9811, p=0.3494, p=0.6500, 2-way ANOVA with SMCT) G- Density of SST cells in each cortical layer in V1, WT v β-KO. The laminar density of SST cells was not changed compared to WT (LtR: p=0.6494, p=0.8858, p=0.9728, p=0.8280, 2-way ANOVA with SMCT)

### The observed adult α-KO phenotype is unlikely to be due to increased repulsion during cIN radial migration or early perinatal cell death

A series of 3 stages of cIN development occur during perinatal time points that could potentially influence cIN spatial distribution: 1) radial migration, which occurs from P2-P5; 2) programmed cell death, which occurs from P4-P12 and peaks at P7 (Southwell DG *et al*. 2012); 3) synaptogenesis, which occurs between P8-P16 (Favuzzi E et al. 2019). Given that the spatial distribution phenotypes observed in α-KO and β-KO were similar but milder in β-KO, we focused on the α-KO for our analysis during perinatal time points: P3, P7 and P13.

Interneuron migration in the context of PV and SST cells is difficult to directly study, due to the relatively late expression of PV (Vogt Weisenhorn DM et al. 1998). Thus, to visualize cells early on, we made use of Nkx2.1^Cre^; Ai9 mice, which labels all MGE derived cINs with tdTomato (Fig4A). This strategy allowed us to study the spatial characteristics of a combined population of PV and SST cells in the context of cPcdh mutations and to infer the timing for when the adult mutant phenotype emerges perinatally.

**Figure 4.**
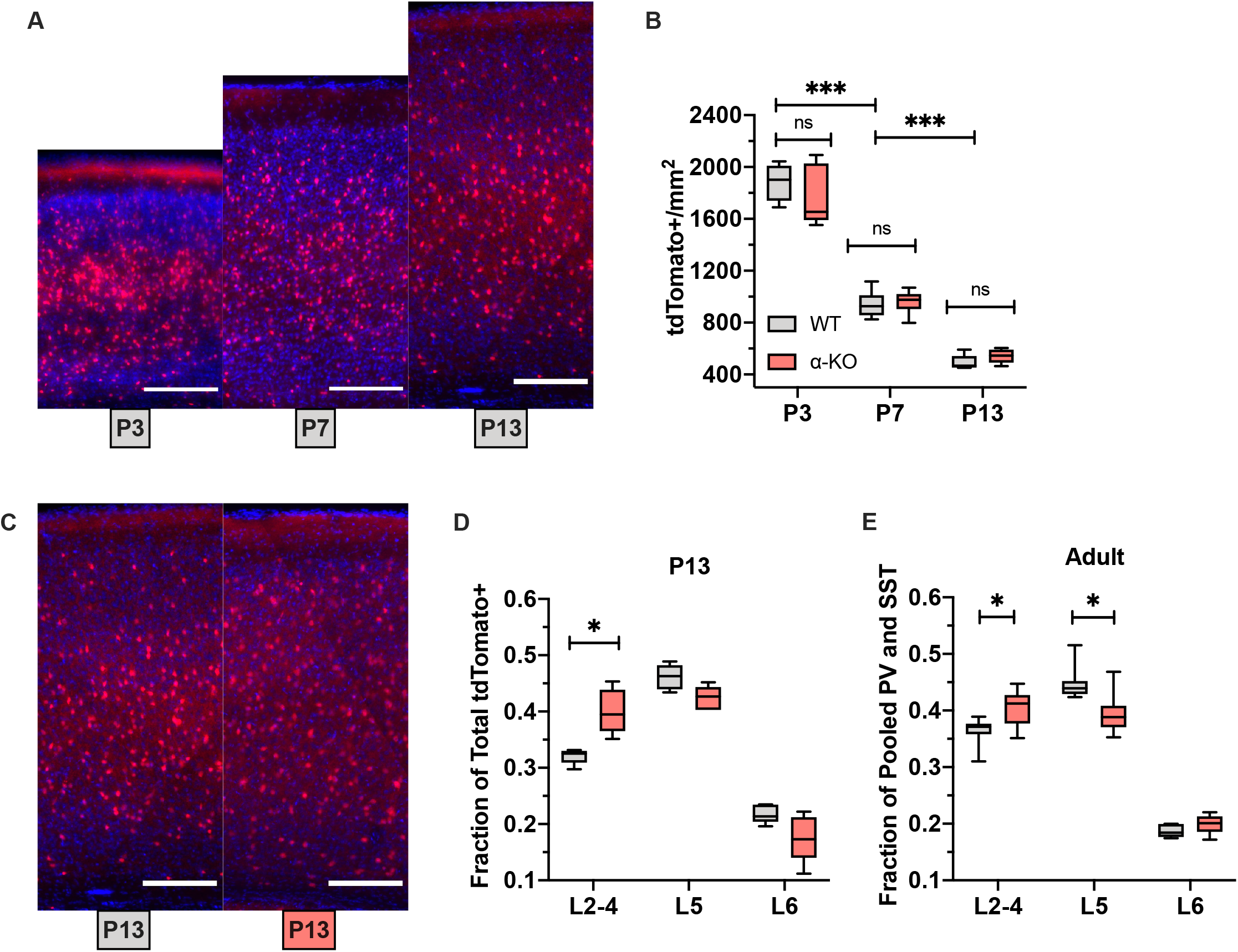
cIN laminar distribution similar to the adult α-KO V1 emerges at postnatal day 13. A-Representative images of tdTomato+ (red) neurons counterstained with DAPI (blue) from Nkx2.1^Cre^;Ai9 V1 at (LtR:) P3, P7, P13; scale bar=200μm B-Density of tdTomato+ cells in V1 at P3, P7, and P13, Nkx2.1^Cre^; Ai9; cPcdh^WT^ vs. α-KO. The density of tdTomato+ cells decreases over time, but does not differ between WT and α-KO (WT v α-KO: P3 p=0.5413, P7 p=0.9899, P13 p=0.9006; p<0.0001 for comparisons between timepoints, 2-way ANOVA with SMCT) C-Representative images of tdTomato+ (red) neurons counterstained with DAPI (blue), from P13 Nkx2.1^Cre^;Ai9;cPcdh^WT^ (left) or Nkx2.1^Cre^;Ai9;α-KO (right) V1; scale bar=200μm D-Relative proportion of tdTomato+ cells in each cortical layer in V1 at P13, Nkx2.1^Cre^; Ai9; cPcdh^WT^ vs. α-KO. The proportion of superficial tdTomato+ cells is increased in the α-KO at P13 (L2-4: p=0.0245, L5: p=0.0779, L6: p=0.2353, 2-way ANOVA with SMCT) E-Relative proportion of pooled PV and SST cells in each cortical layer from V1 P30 α-KO. Proportion in superficial layers is increased, while L5 is decreased, similar to the P13 Nkx2.1^Cre^;Ai9;α-KO (L2-4: p=0.0282, L5: p=0.0129, L6: p=0.1683, 2-way ANOVA with SMCT)

Consistent with prior reports (Southwell DG *et al*. 2012), the density of tdTomato+ cells decreases between P3 and P13 (Fig4B), with no differences between WT and α-KO (Supp7A). This coincides with an expansion in cortical thickness during the same time frame (Supp7A). At P3 and P7, we observe no changes in the relative laminar distribution of tdTomato+ cells (Supp7C, D). However, at P13, there is a significant increase in the fraction of tdTomato+ cells in upper layers (Fig4C, D) and a trend towards reduced L5 (p=0.0779). In the perinatal datasets, it was difficult to distinguish L4 from L2/3 using DAPI, but the border between L4 and L5 was always clear. In order to compare our results to the adult α-KO results, we pooled PV and SST, as they encompass the entire Nkx2.1 lineage in the cortex (Rudy B *et al*. 2011), and also pooled L2/3 and L4. Here, we found that the laminar distribution in the adult α-KO was similar to that of the P13 Nkx2.1Cre;Ai9;α-KO (Fig4E).

### Loss of a single protocadherin, pcdh-αc2, alters the laminar distribution of SST cells but does not change cell density

Our data indicate that the loss of cPcdhs differentially affects PV and SST neurons, yet the majority of cPcdh isoforms are expressed stochastically in most if not all cortical neurons (Chen WV and T Maniatis 2013). The exceptions are pcdh-αc1, αc2, γc3, γc4, and γc5, which are transcriptionally-regulated (Kawaguchi M et al. 2008; Chen WV and T Maniatis 2013). Although these regulated isoforms are broadly expressed, single cell RNA sequencing data from the Allen Brain Institute (Tasic B et al. 2018) suggests that certain classes of cells have enriched expression. Pcdh-αc2 is highly expressed in PV cells, and to a lesser extent SST, in adult V1 (Supp 8). Based on this data, we explored the contribution of pcdh-αc2 to the α-KO phenotype using a pcdh-αc2 specific knockout mouse (αc2-KO).

While pcdh-αc2 is enriched in adult PV cells, we did not observe significant changes to PV density or NND in αc2-KO mice (Fig5A, B). Laminar distribution was also unchanged for PV cells (Fig5C). With SST cells, however, we found that the αc2-KO recapitulated some aspects of the α-KO phenotype: a significant increase in the relative amount of SST cells in L2/3 (Fig5D) without a change in SST density or NND (Fig5A,B). Unlike the α-KO we did not see a significant reduction in L5 (Fig5D). We also did not observe a change in cortical thickness (Supp9A-C) or CV (Supp9D) in the αc2-KO. Together this data suggests that pcdh-αc2 may be playing a larger role in SST cell layer distribution compared to PV.

**Figure 5.**
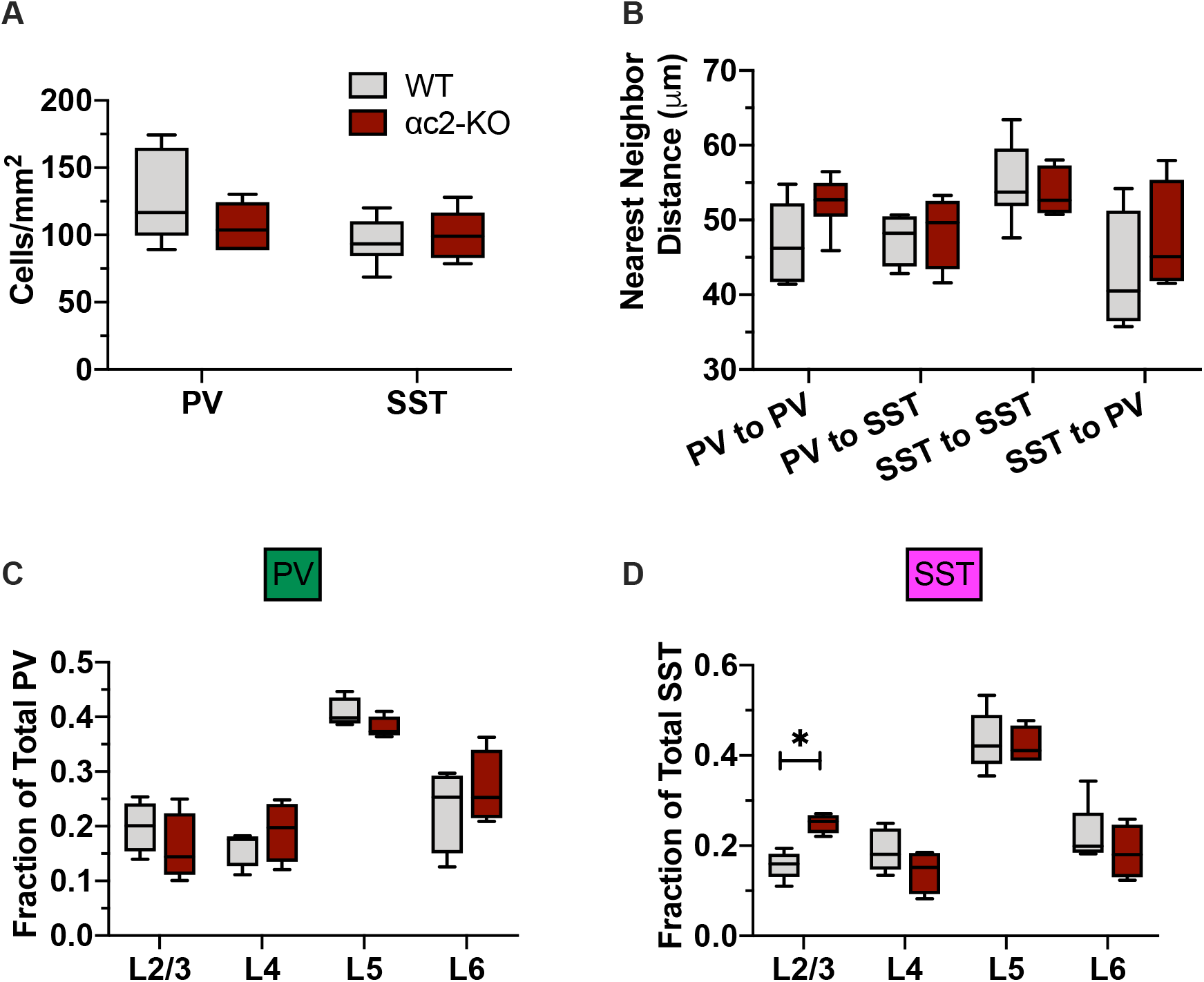
Loss of a single protocadherin, pcdh-αc2, alters the laminar distribution of SST cells but does not change cell density. A-Density of PV and SST cells in V1, WT vs. αc2-KO. Cell density is not changed in the αc2-KO (PV: p=0.2310; SST: p=0.9153, 2-way ANOVA with SMCT) B-NND between PV-PV, PV-SST, SST-SST, and SST-PV pairs. NND is not changed in the αc2-KO between any group compared (LtR: p=0.3259, p>0.9999, p=0.9667, p=0.7689, 2-way ANOVA with SMCT) C-Relative proportion of PV cells in each cortical layer in V1, WT vs. αc2-KO. The relative proportion of PV cells was not changed (LtR: p=0.8338, p=0.8728, p=0.5121, p=0.9374, 2-way ANOVA with SMCT) D-Relative proportion of SST cells in each cortical layer in V1, WT vs. αc2-KO. There are relatively more SST cells in L2/3 in the αc2-KO compared to WT but other layers are not altered (LtR: p=0.0441, p=0.5573, p=0.9974, p=0.7552, 2-way ANOVA with SMCT) E-Density of PV cells in each cortical layer in V1, WT vs. αc2-KO. PV density is not significantly changed in any layer in the αc2-KO, though L5 does trend lower (LtR: p=0.6515, 0.7319, 0.1744, 0.9101, 2-way ANOVA with SMCT) F- Density of SST cells in each cortical layer in V1, WT vs. αc2-KO. SST density is not significantly changed in any layer in the αc2-KO, though it does trend up in L2/3 (LtR: p=0.2141, p=0.3664, p=0.7561, p=0.9115, 2-way ANOVA with SMCT)

## Discussion

The spatial positioning of neurons can be critical to their function, as is the case in sensory neurons in *Drosophila* (Matthews BJ et al. 2007; Liu C et al. 2020) or starburst amacrine cells in retina (Fuerst PG *et al*. 2008; Garrett AM et al. 2018; Ing-Esteves S et al. 2018). In this study, we analyzed the spatial position of cINs in WT and cPcdh mutants. cINs originate from distal progenitor zones and distribute throughout the cortex during development through extensive migration and cell death (Tremblay R *et al*. 2016; Wamsley B and G Fishell 2017). We hypothesized that cIN subclasses self-recognize during development to ensure that subclass-specific circuits are evenly distributed. We also considered the possibility that generic repulsion might exist between all cINs to further facilitate even distribution of inhibitory networks. In our detailed analysis of cIN spatial distribution in the WT however, PV and SST cells appear to ignore the presence of the other cell type even though both cells occupy similar laminar domains. This suggests that generic interneuron-interneuron repulsion does not occur. Rather, our data indicate that cell-cell repulsion only exists within cIN subclasses and not between them. This is reminiscent of the retina in which starburst amacrine cell subtypes are patterned independently of heterotypic cells but are mostly non-randomly spaced with respect to homotypic cells (Fuerst PG *et al*. 2008).

Our examination of cPcdh mutants revealed cIN subclass specific alterations in spatial distribution, cell density and laminar distribution. For instance, we observed that PV density was significantly reduced in the α-KO, while SST density was unchanged. PV density reduction was accompanied by a significant increase in PV cell NND. A recent study reported that the γ-cluster (but not α- or β-clusters) is important for cIN survival (Mancia Leon WR *et al*. 2020). It is important to note that in this study, the authors quantified Nkx2.1^Cre^ fate mapped neurons without distinguishing between PV and SST in their analysis of α-KO and β-KO mice. Thus, the decrease in PV density we observed may have been masked by quantifying the entire Nkx2.1 (MGE) lineage as whole. Interestingly, however, Mancia and colleagues note trends towards decreased Nkx2.1 lineage cell density, consistent with our findings. Further, in the spinal cord γ-cluster cPcdhs generally mediate interneuron survival, but FoxP2+ subtypes of spinal cord interneurons are reduced in both α- and β-cluster mutants (Hasegawa S *et al*. 2016). While the mechanism driving the decrease in PV density we observe is unclear, α-KO PV cells exhibit connectivity defects including reduced inhibitory synapses, reduced neurite number, and reduced neurite length (Shao Z et al. 2019). As such, alterations in connectivity may underlie increased PV cell death, resulting in decreased cell density. We also observed subtype specific changes in the relative laminar distribution of V1 α-KO cINs. Both PV and SST cells were reduced in L5, but PV cells were increased in L6 while SST cells were increased in L2/3. We noted similar shifts in laminar distribution in S1 and in the β-KO, although some of these effects were trends rather than significant changes.

While we observed clear subtype specific phenotypes in laminar distribution and cell density in cPcdh mutants, most cPcdhs isoforms are stochastically expressed (Chen WV and T Maniatis 2013). To reconcile this, we examined existing single cell RNA sequencing datasets of adult mouse visual cortex (Tasic B *et al*. 2018) and found that a non-stochastic member (Kawaguchi M *et al*. 2008; Chen WV and T Maniatis 2013) of the α-cPcdhs, pcdh-αc2, was highly expressed in subgroups of PV and SST cells. When we examined spatial distribution of PV and SST in αc2-KO mice, we found that L2/3 SST cells were increased, while SST cells in other layers and all PV cells were mostly unaffected. This finding supports the notion that non-stochastic subtype specific expression of cPcdhs helps regulate cIN distribution. While it is not clear why SST cells are specifically affected, it is possible that pcdh-αc2 expression is enriched early in SST cells, whereas enriched PV expression occurs after the mutant phenotype emerges (P13).

Taken together, we observed subclass-specific laminar distribution phenotypes that were strikingly similar across three different cPcdh mutants and in two different cortical areas (V1 and S1). Loss of cPcdh diversity can aberrantly increase recognition and repulsion between neurons that don’t normally repel (Lefebvre JL *et al*. 2012). Further, shortly after birth, interneurons undergo radial migration into the cortical plate to settle in their final layer (Marin O 2013). Thus, an appealing model is that aberrant cPcdh matching due to decreased cPcdh diversity would in turn alter subclass spacing. Here, cIN radial migration would be an ideal process for cPcdh regulation since laminar position is largely determined in this phase. However, when we examined α-KO mice during perinatal time points, we did not observe changes in laminar distribution until P13, well after radial migration has ceased.

The precise mechanism for how loss of cPcdh diversity alters cIN subclass spacing remains to be determined. One possibility is cell intrinsic: Previous studies have found that α-KO PV cells possess morphological defects and reduced inhibitory synapse density, while excitatory synapse density was not changed (Shao Z *et al*. 2019), suggesting defects in intrinsic connectivity. Another possibility, perhaps acting in parallel, is that loss of cPcdh diversity increases the likelihood of neighboring cINs to mistakenly identifying non-self neurites as self neurites, as has been observed in retina (Lefebvre JL *et al*. 2012). In the cortex, PV and SST cells occupy similar spatial domains and densely populate deep layers, in particular L5. cPcdh diversity could therefore enable cINs of the same subclass to pack into high density areas such as deep layers of the cortex. cIN target innervation is similarly dense, with many cINs targeting the same pyramidal cell, and many pyramidal cells being inhibited by the same cIN (Fino E and R Yuste 2011; Packer AM *et al*. 2013). Both cell intrinsic and extrinsic defects could then lead to altered connectivity and reduced activity, which would reduce the number of cINs (Priya R et al. 2018) in specific areas, and most prominently in areas where cINs are most dense. This would potentially account for why the phenotype in L5 of V1 was more prominent - V1 is significantly thinner than S1, which would place even more spatial constraint on cINs and increase the chances of encountering an aberrantly matched neuron.

Alternatively, alterations in cIN laminar distribution could be due to selective survival during development as a result of alterations is cortical activity. cIN survival is regulated by neuronal activity (Priya R *et al*. 2018; Wong FK et al. 2018), and aberrant neuronal activity can alter the laminar positioning in cINs of the caudal ganglionic eminence (De Marco Garcia NV et al. 2011). Deficits in serotonergic wiring have also been reported in the α-KO and αc1/2-KO mice (Katori S et al. 2009; Chen WV *et al*. 2017) and alterations in serotonergic tone could also affect overall interneuron activity. Interestingly, deficits in visual acuity and aggregation of retinogeniculate terminals have been reported to appear between P10 and P14 in the α-KO (Meguro R et al. 2015), coinciding with the emergence of cIN phenotype we observed at P13. This could alter the activity state of the visual cortex, where we observe the most robust mutant phenotype.

Cortical interneurons function by establishing a distributed network of locally-projecting neurons throughout the cortex. Our study indicates that while cPcdhs do not affect subclass cell-cell repulsion, they play an important role in regulating how cINs spatially distribute among cortical layers. The findings also demonstrate that cPcdh loss-of-function differentially affects interneuron subtypes. Indeed, specific genetic ablation of αc2 largely phenocopies the mutant SST phenotype of the whole cluster a-KO, while PV cells are unaffected. This suggests that non-stochastic cPcdh expression helps establish subtype-specific spatial distribution patterns, possibly through subtype-specific expression. Finally, when we profiled cIN spatial arrangement during perinatal timepoints, we found that the α-KO phenotype only emerged later at P13, well after radial migration, and coinciding with cIN morphological elaboration and synaptogenesis. Thus, our work raises the intriguing possibility that cPcdh isoforms, expressed with subtype specificity, govern early circuit integration and thereby differential cIN cell survival. It remains to be determined whether this is the result of cell-autonomous or circuit-level mechanism; Previous findings suggest both could be at play, and that both could serve as compelling avenues to pursue for future studies.

## Acknowledgements

The authors would like to thank Tom Maniatis for generously providing cPcdh mutant animals, as well as Chiamaka Nwakeze and Amy Kirner for their help with these animals. Thanks to Luke Hammond and Tu Nguyen for their help with imaging, and Kevin Sitek for help with coding. Also thanks to Wes Grueber, Carol Mason, Franck Polleux, Richard Vallee and Hynek Wicherle for thoughtful comments on the project and the manuscript. Finally thank you to Luke Nunnelly, Patrick Dummer, Sofia Leal Santos and Melissa McKenzie for their feedback and discussion. This work was supported by the Irma T. Hirschl Foundation and R03MH119443 to EA.

## Glossary

ABAN: Allen Brain Atlas Number
CV: Coefficient of variance of the nearest neighbor distance
DMCT: Dunnett’s multiple comparisons test
L2/3, L4, L5, L6: Cortical layers 2/3, 4, 5, 6
LtR: Left to right
NND: Nearest neighbor distance
P3, P7, P13: Post-natal day 3, 7, 13
PCF: Paired correlation function
PV: Parvalbumin
S1: Primary somatosensory cortex
SMCT: Sidak’s multiple comparisons test
SST: Somatostatin
TMCT: Tukey’s multiple comparisons test
V1: Primary visual cortex
WT: Wild Type
α-KO: Alpha clustered protocadherin knockout mouse
αc2-KO: Protocadherin αc2 knockout mouse
β-KO: Beta clustered protocadherin knockout mouse

## Supplemental Figure Legends

**Supplemental Figure 1.**
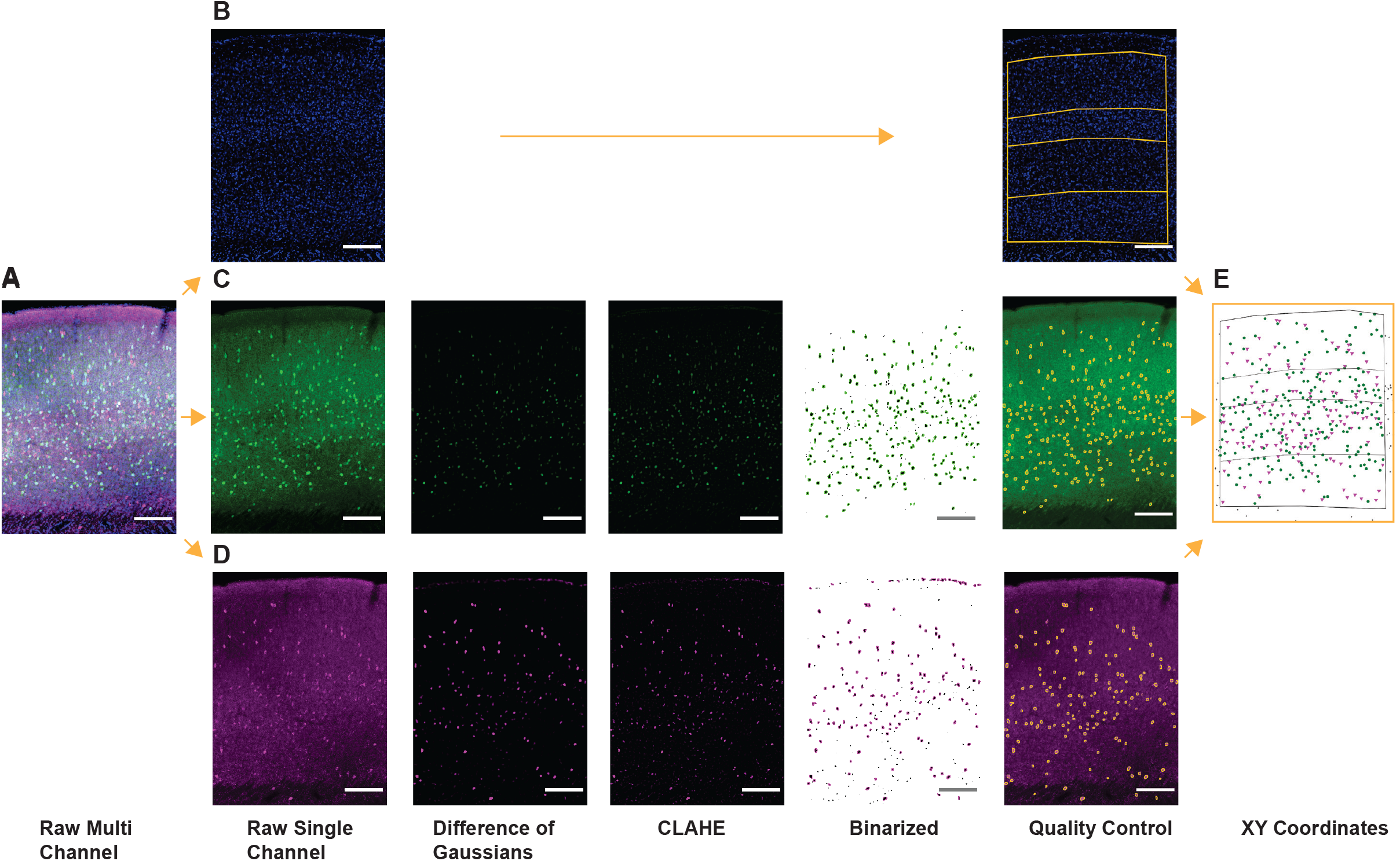
Semi-automated image quantification pipeline for spatial analysis of cINs. A-Representative fluorescent image of WT V1 labeled with DAPI (blue), anti-PV (green), and anti-SST (magenta); scale bar=200μm B-Workflow for DAPI-containing channel. L: Representative fluorescent image of DAPI. R: The density of DAPI stained nuclei is used to manually draw layers onto the image. The XY coordinates of these layers are saved. C-Workflow for PV-containing channel. LtR: Original image, image after difference of Gaussians, image after CLAHE, binarized image with outlines of size and circularity gated particles, final cell count after manual quality control overlaid on cell image D-Workflow for SST-containing channel. LtR: Original image, image after difference of Gaussians, image after CLAHE, binarized image with outlines of size and circularity gated particles, final cell count after manual quality control overlaid on cell image E-Final image generated in Spatstat using XY coordinates obtained from images

**Supplemental Figure 2.**
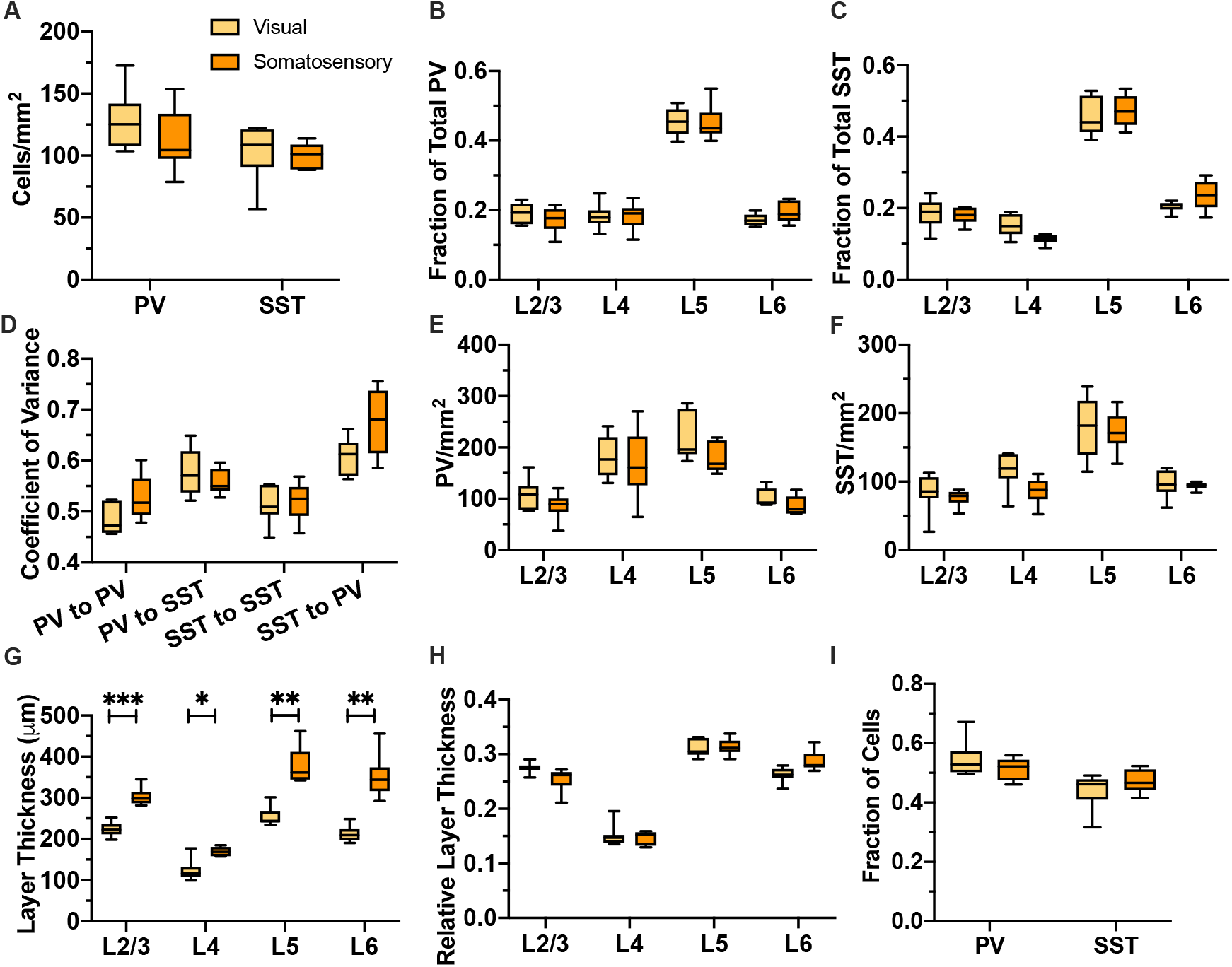
Measurements from S1 are not significantly different from V1 except for cortical thickness. A-PV and SST cell density, WT V1 vs. S1. cIN density does not significantly differ between V1 (yellow) and S1 (orange) cortex (PV: p=0.3453, SST: p =0.9726, 2-way ANOVA with SMCT) B-Relative proportion of PV cells in each layer, WT V1 vs. S1. The laminar distribution of PV cells was not different between V1 and S1 (LtR: p=0.7889, >0.999, >0.999, 0.7327, 2-way ANOVA with SMCT) C-Relative proportion of SST cells in each layer, WT V1 vs. S1. The laminar distribution of SST cells was not different between V1 and S1 (LtR: p=0.9939, p=0.2343, p=0.9416, p=0.3985, 2-way ANOVA with SMCT) D-CV between PV-PV, PV-SST, SST-SST, SST-PV pairs, WT V1 vs. S1. The CV was not different between V1 and S1 (LtR: p=0.2394, p=0.8499, p=0.9993, p=0.2153, 2-way ANOVA with SMCT) E-Density of PV cells in each layer, WT V1 vs. S1. PV density was not different in any layer between V1 and S1 (LtR: p=0.6495, p=0.9957, p=0.3367, p=0.4709, 2-way ANOVA with SMCT) F-Density of SST cells in each layer, WT V1 vs. S1. SST density was not different in any layer between V1 and S1 (LtR: p=0.9689, p=0.1389, p=0.9998, p=0.9991, 2-way ANOVA with SMCT) G-Cortical thickness, WT V1 vs. S1. Each cortical layer was significantly thicker in S1 compared to V1 (LtR: p=0.0002, p=0.0104, p=0.0019, p=0.0045, 2-way ANOVA with SMCT) H-Relative cortical thickness, WT V1 vs. S1. Proportionally, cortical layers were similar in V1 and S1 (LtR: p=0.2508, p=0.9760, p=0.9989, p=0.0978, 2-way ANOVA with SMCT) I-Fraction of all cells that were PV or SST, WT V1 vs. S1. The fraction of all cells that were PV or SST was not different between V1 and S1 (PV: p=0.4455, SST: p=0.4463, 2-way ANOVA with SMCT)

**Supplemental Figure 3.**
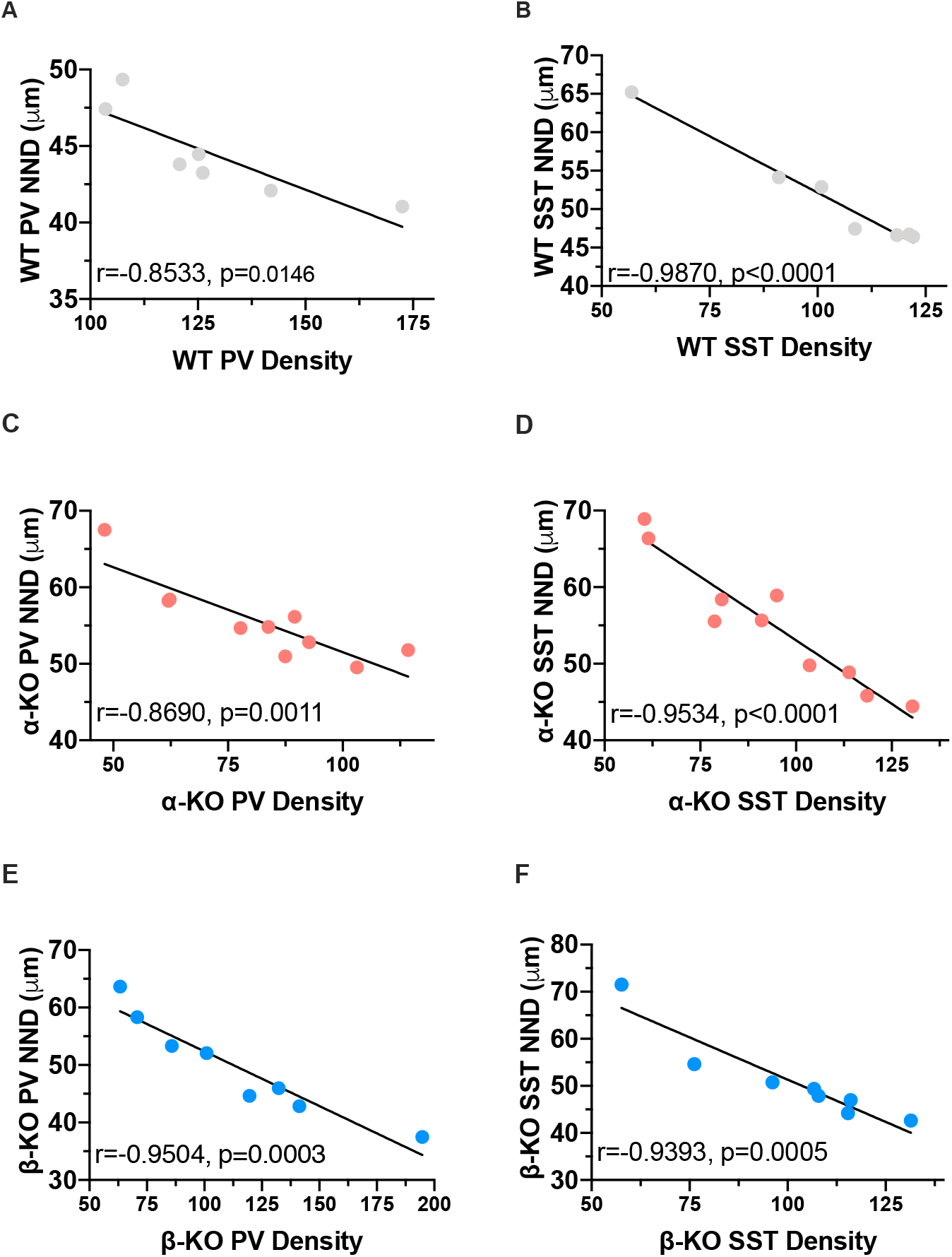
Nearest neighbor distance is significantly negatively correlated with cell density. A-Correlation between WT PV-PV NND and PV density, V1. WT PV NND and PV density are significantly negatively correlated (r=-0.8533, p=0.0146, simple linear regression) B-Correlation between WT SST-SST NND and SST density, V1. WT SST NND and SST density are significantly negatively correlated (r=-0.9870, p<0.0001, simple linear regression) C-Correlation between α-KO PV-PV NND and PV density, V1. α-KO PV NND and PV density are significantly negatively correlated (r=-0.8690, p=0.0011, simple linear regression) D-Correlation between α-KO SST-SST NND and SST density, V1. α-KO SST NND and SST density are significantly negatively correlated (r=-0.9534, p<0.0001, simple linear regression) E-Correlation between β-KO PV-PV NND and PV density, V1. β-KO PV NND and PV density are significantly negatively correlated (r=-0.9504, p=0.0003, simple linear regression) F-Correlation between β-KO SST-SST NND and SST density, V1. β-KO SST NND and SST density are significantly negatively correlated (r=-0.9393, p=0.0005, simple linear regression)

**Supplemental Figure 4.**
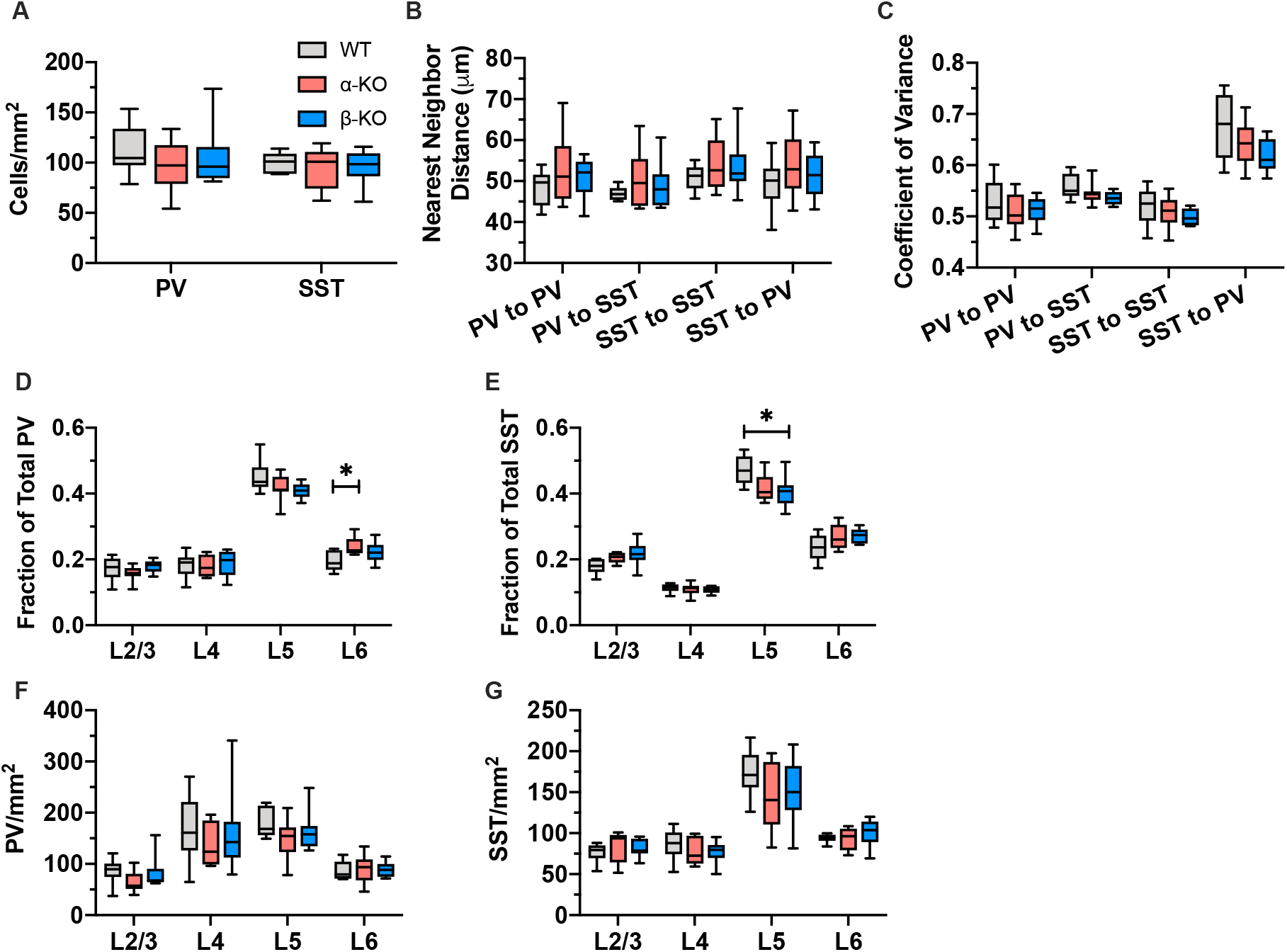
Loss of cPcdh diversity alters laminar distribution in S1. A-Density of PV and SST cells in S1, WT vs. α-KO v β-KO. Cell density is not altered in S1 (PV: WT v α-KO p=0.3796, WT v β-KO p=0.8432; SST: WT v α-KO p=0.8511, WT v β-KO p=0.9192, 2-way ANOVA w Tukey’s MCT) B-NND between PV-PV, PV-SST, SST-SST, SST-PV pairs in S1, WT vs. α-KO vs. β-KO. Nearest neighbor distance is not altered in S1 (PV-PV: WT v α-KO p=0.3563, WT v β-KO p=0.6044; PV-SST: WT v α-KO p=0.3440, WT v β-KO p =0.5523; SST-SST: WT v α-KO p=0.4379, WT v β-KO p=0.5069; SST-PV: WT v α-KO p=0.4596, WT v β-KO p=0.8517, 2-way ANOVA w TMCT) C-CV between PV-PV, PV-SST, SST-SST, SST-PV pairs in S1, WT vs. α-KO vs. β-KO. CV is not altered in S1 (PV-PV: WT v α-KO p=0.6905, WT v β-KO p=0.7404; PV-SST: WT v α-KO p=0.5066, WT v β-KO p =0.1866; SST-SST: WT v α-KO p=0.8681, WT v β-KO p=0.4300; SST-PV: WT v α-KO p=0.5404, WT v β-KO p=0.1943, 2-way ANOVA w TMCT) D-Relative proportion of PV cells in each cortical layer in S1, WT vs. α-KO vs. β-KO. The relative amount of PV cells was increased in L6 in the α-KO. (L2/3: WT v α-KO p=0.6714, WT v β-KO p=0.9203; L4: WT v α-KO p=0.9951, WT v β-KO p =0.9256; L5: WT v α-KO p=0.4976, WT v β-KO p=0.2247; L6: WT v α-KO p=0.0326, WT v β-KO p=0.2465, 2-way ANOVA w TMCT) E- Relative proportion of SST cells in each cortical layer in S1, WT vs. α-KO vs. β-KO. The relative amount of SST cells was decreased in L5 of the β-KO. SST trended up in L2/3 in α-KO and β-KO, but these p values were barely above the statistical significance threshold. Similarly, L5 of α-KO trended lower. (L2/3: WT v α-KO p=0.0894, WT v β-KO p=0.0613; L4: WT v α-KO p=0.8437, WT v β-KO p=0.6363; L5: WT v α-KO p=0.0820, WT v β-KO p=0.0363; L6: WT v α-KO p=0.3088, WT v β-KO p=0.2059, 2-way ANOVA w TMCT) F-Density of PV cells in each cortical layer in S1, WT vs. α-KO vs. β-KO. PV density was not different across layers (L2/3: WT v α-KO p=0.2578, WT v β-KO p=0.9347; L4: WT v α-KO p=0.6097, WT v β-KO p=0.9825; L5: WT v α-KO p=0.1882, WT v β-KO p=0.6188; L6: WT v α-KO p=0.9735, WT v β-KO p=0.9663, 2-way ANOVA w TMCT) G-Density of SST cells in each cortical layer in S1, WT vs. α-KO vs. β-KO. SST density was not different across layers (L2/3: WT v α-KO p=0.7389, WT v β-KO p=0.6128; L4: WT v α-KO p=0.7432, WT v β-KO p=0.5920; L5: WT v α-KO p=0.2925, WT v β-KO p=0.4476; L6: WT v α-KO p=0.9864, WT v β-KO p=0.5703, 2-way ANOVA w TMCT)

**Supplemental Figure 5.**
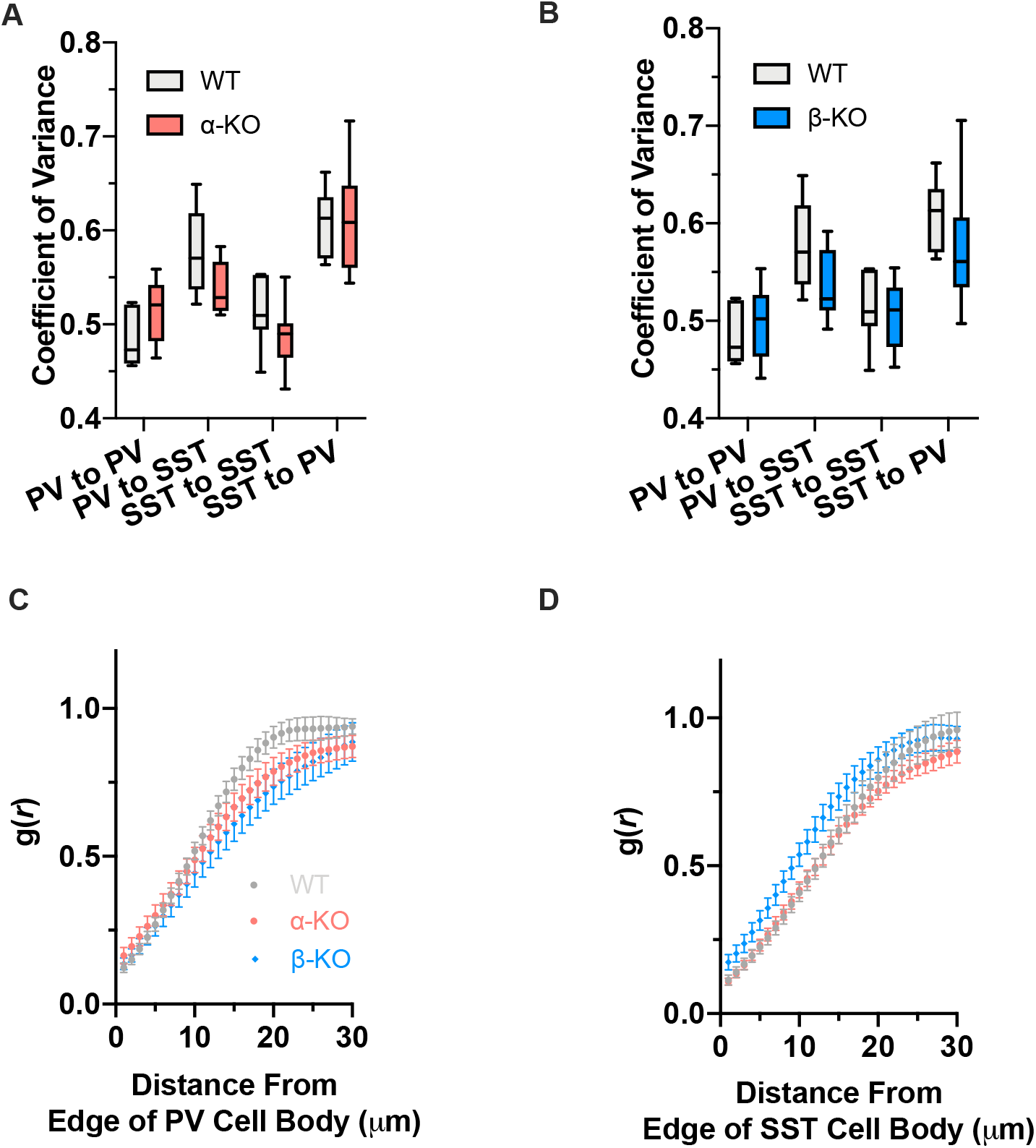
Regularity of spacing is not altered by loss of cPcdh diversity. A-CV between PV-PV, PV-SST, SST-SST, SST-PV pairs in V1, WT vs. α-KO. CV is not significantly different between WT and α-KO (LtR: p=0.3317, p=0.1717, p=0.4059, p>0.9999, 2-way ANOVA with SMCT) B-CV between PV-PV, PV-SST, SST-SST, SST-PV pairs in V1, WT vs. β-KO. CV is not significantly different between WT and β-KO (LtR: p=0.8832, p=0.2590, p=0.9781, 0.5915, 2-way ANOVA with SMCT) C-PCF of PV-PV pairs in V1, WT vs. α-KO vs. β-KO (No p<0.05 for any distance, 2-way ANOVA with SMCT) D-PCF of SST-SST pairs in V1, WT vs. α-KO vs. β-KO (No p<0.05 for any distance, 2-way ANOVA with SMCT)

**Supplemental Figure 6.**
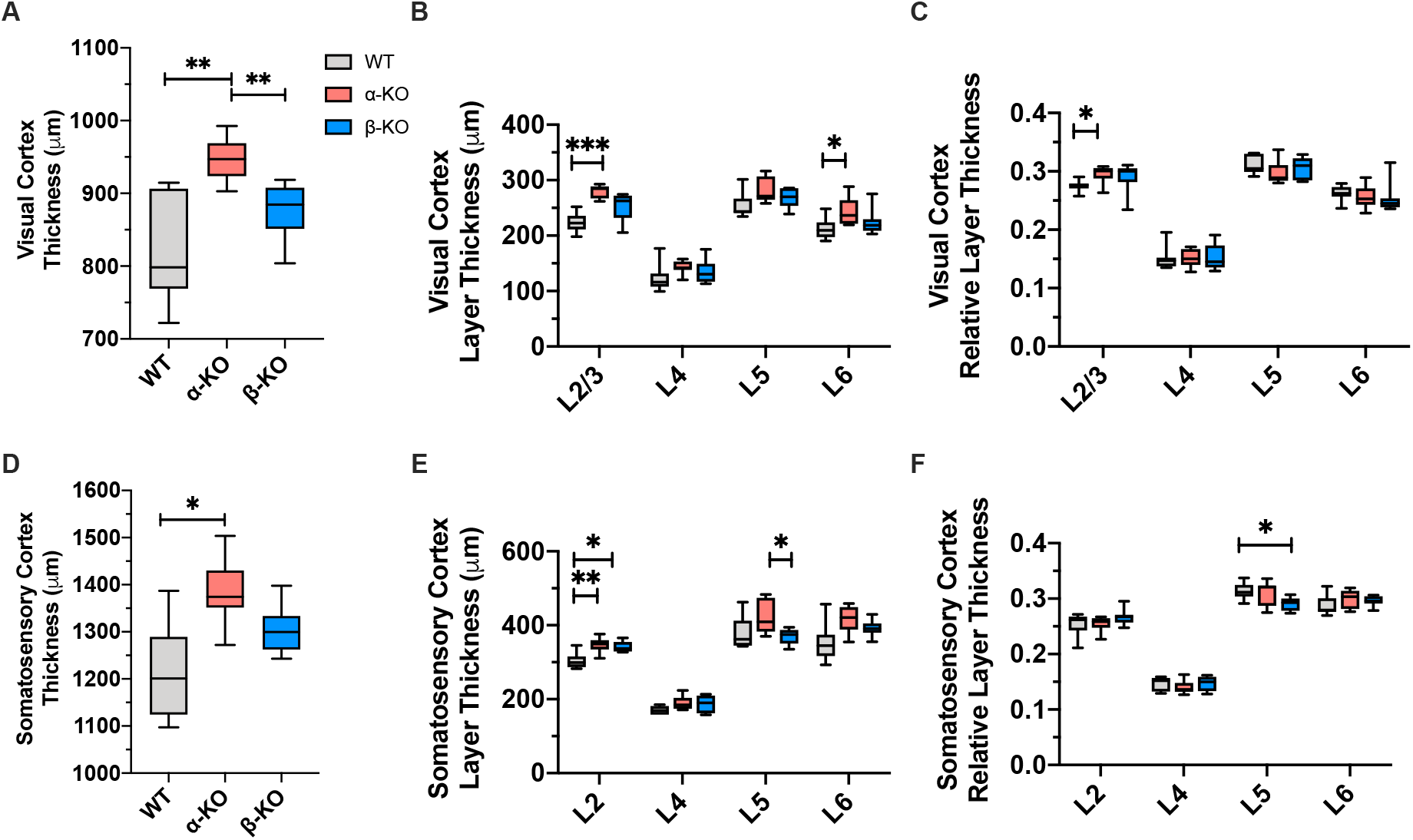
The α-KO is thicker in both V1 and S1 compared to WT. A-Thickness of V1, WT v α-KO vs. β-KO. The α-KO cortex is thicker than WT cortex in V1 (WT v α-KO p=0.0066, WT v β-KO p=0.1711, 1-way ANOVA with DMCT) B-Laminar thickness of V1, WT vs. α-KO vs. β-KO.In V1, L2/3 and L6 of the α-KO cortex is thicker than WT (L2/3: WT v α-KO p=0.0001, WT v β-KO p=0.0550; L4: WT v α-KO p=0.2183, WT v β-KO p=0.6690; L5: WT v α-KO p=0.0689, WT v β-KO p=0.4240; L6: WT v α-KO p=0.0305, WT v β-KO p=0.6025, 2-way ANOVA w TMCT) C-Relative laminar thickness of V1, WT v α-KO v β-KO. L2/3 in the α-KO is increased relative to other layers but other layers are proportionally unaltered (L2/3: WT v α-KO p=0.0154, WT v β-KO p=0.3006; L4: WT v α-KO p=0.9990, WT v β-KO p=0.9883; L5: WT v α-KO p=0.2493, WT v β-KO p=0.7473; L6: WT v α-KO p=0.7295, WT v β-KO p=0.6692, 2-way ANOVA w TMCT) D-Thickness of S1, WT v α-KO vs. β-KO. The α-KO cortex is thicker than WT cortex in S1 (WT v α-KO p=0.0163, WT v β-KO p=0.2243, 1-way ANOVA with DMCT) E-Laminar thickness of S1, WT vs. α-KO vs. β-KO. In S1, L2/3 is thicker in both the α-KO and β-KO. L5 of the α-KO was thicker than β-KO but not statistically significant compared to WT. (L2/3: WT v α-KO p=0.0073, WT v β-KO p=0.0159; L4: WT v α-KO p=0.0517, WT v β-KO p=0.1467; L5: WT v α-KO p=0.1791, WT v β-KO p=0.9178; L6: WT v α-KO p=0.0774, WT v β-KO p=0.2747, 2-way ANOVA w TMCT) F-Relative laminar thickness of V1, WT vs. α-KO vs. β-KO. Proportionally, most layers are unchanged compared to WT, except L5 in the β-KO which is reduced (L2/3: WT v α-KO p=0.9985, WT v β-KO p=0.4695; L4: WT v α-KO p=0.5299, WT v β-KO p>0.9999; L5: WT v α-KO p=0.8217, WT v β-KO p=0.0346; L6: WT v α-KO p=0.3954, WT v β-KO p=0.5858, 2-way ANOVA w TMCT)

**Supplemental Figure 7.**
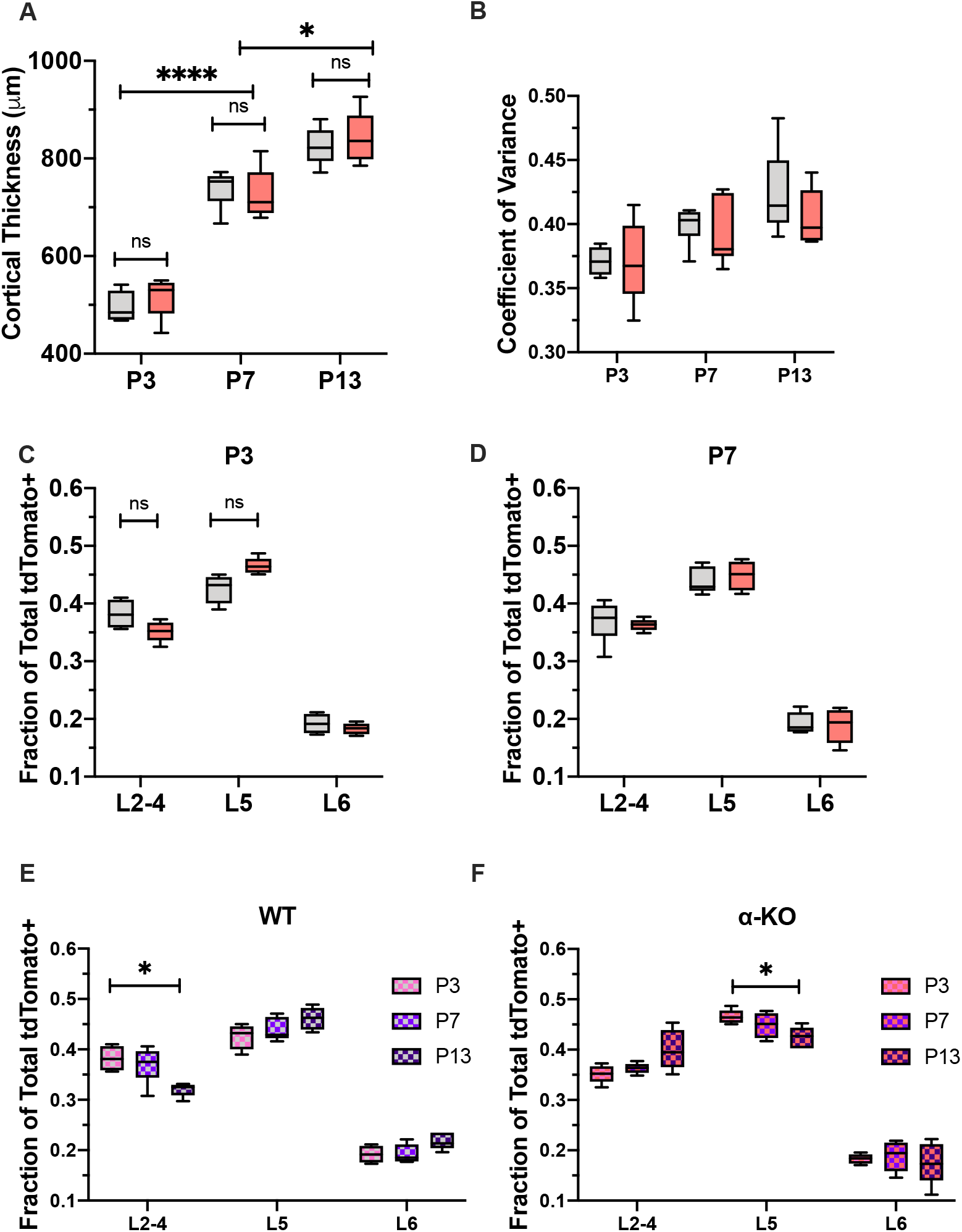
Laminar distribution is not altered at P3 or P7, mutant phenotype emerges at P13. A-Thickness of V1 at P3, P7, and P13, Nkx2.1^Cre^; Ai9; cPcdh^WT^ vs. α-KO. Cortical thickness increases over time but is not different between WT and α-KO (WT v α-KO: P3 p=0.7857, P7 p=0.9623, P13 p=0.9341; WT P3 v P7: p<0.0001 WT P7 v P13: p=0.0101, 2-way ANOVA with SMCT) B- CV between tdTomato+ cells in V1 at P3, P7, and P13, Nkx2.1^Cre^; Ai9; cPcdh^WT^ vs. α-KO. CV is not different between WT and α-KO at any timepoint (LtR: p=0.9962, p=0.8579, 2-way ANOVA with SMCT) C-Relative proportion of tdTomato+ cells in each layer of V1 at P3, Nkx2.1^Cre^; Ai9; cPcdh^WT^ vs. α-KO. The relative distribution of tdTomato+ cells is not different at P3 (LtR: p=0.2558, p=0.1327, p=0.7794, 2-way ANOVA with SMCT) D-Relative proportion of tdTomato+ cells in each layer of V1 at P7, Nkx2.1^Cre^; Ai9; cPcdh^WT^ vs. α-KO. The relative distribution of tdTomato+ cells is not different at P7 (LtR: p=0.9754, p=0.8558, p=0.9885, 2-way ANOVA with SMCT) E-Relative proportion of tdTomato+ cells in each layer of Nkx2.1^Cre^; Ai9; cPcdh^WT^ V1 at P3, P7, P13. In the WT, the relative amount of tdTomato+ cells in L2/3 decreases over time (L2-4: P3 v P13 p=0.0321, P3 v P7 p=0.8965, P7 v P13 p =0.0672; L5: P3 v P13 p=0.2353, P3 v P7 p=0.8564, P7 v P13 p=0.3917; L6: P3 v P13 p=0.1882, P3 v P7 p>0.9999, P7 v P13 p=0.1385, 2-way ANOVA with TMCT) F- Relative proportion of tdTomato+ cells in each layer of Nkx2.1^Cre^; Ai9; α-KO V1 at P3, P7, P13. In the α-KO, the relative amount of tdTomato+ cells in L5 decreases over time (L2-4: P3 v P13 p=0.1620, P3 v P7 p=0.6177, P7 v P13 p =0.2926; L5: P3 v P13 p=0.0331, P3 v P7 p=0.5278, P7 v P13 p=0.3372; L6: P3 v P13 p=0.9789, P3 v P7 p=0.9744, P7 v P13 p=0.9387, 2-way ANOVA with TMCT)

**Supplemental Figure 8.**
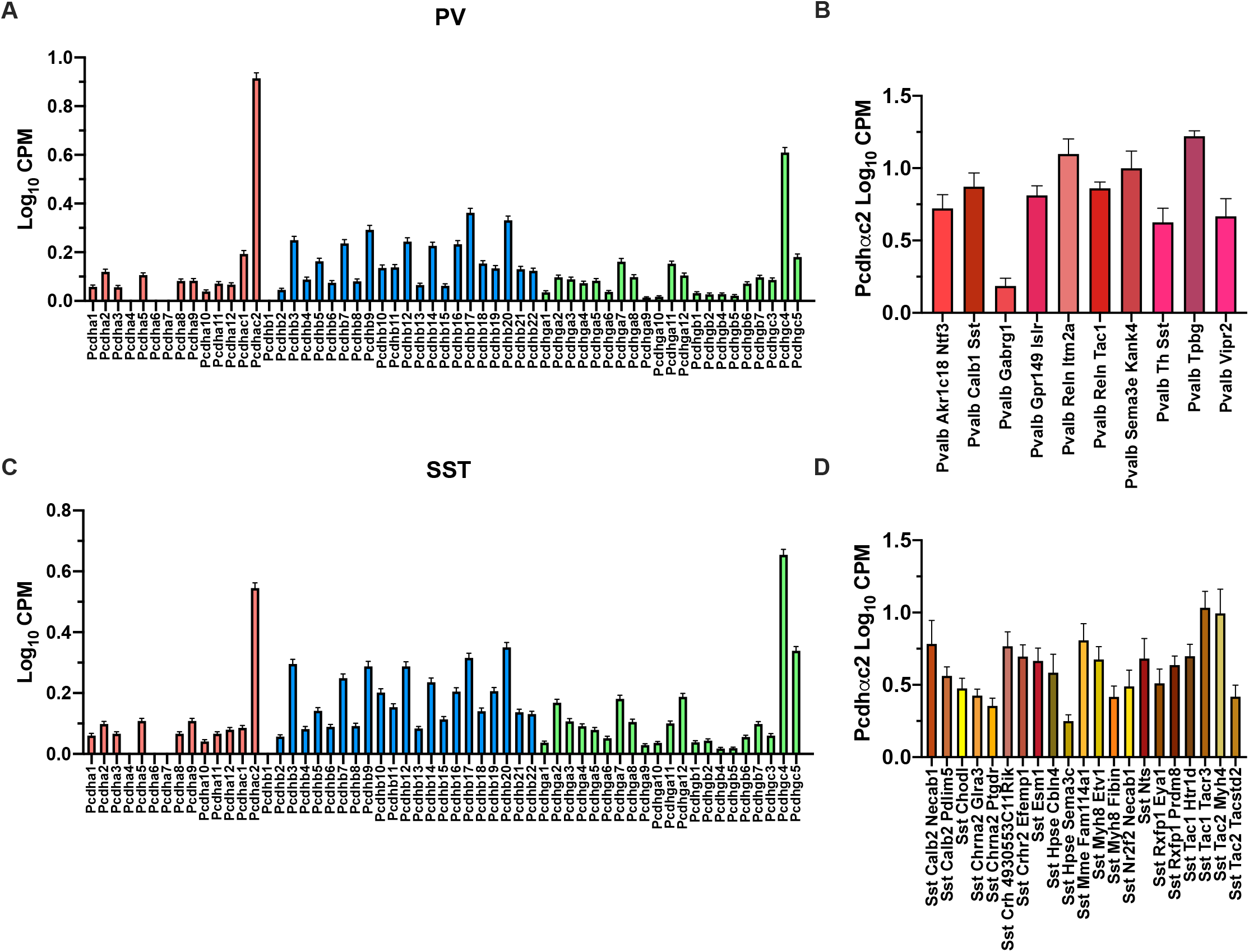
Pcdh-αc2 is expressed at relatively high levels in PV and SST subtypes. A-cPcdh RNA expression in PV cells dissected from adult V1 B-cPcdh RNA expression in SST cells dissected from adult V1 C-Pcdh-αc2 expression in PV subtypes defined by Tasic 2018 D-Pcdh-αc2 expression in SST subtypes defined by Tasic 2018

**Supplemental Figure 9.**
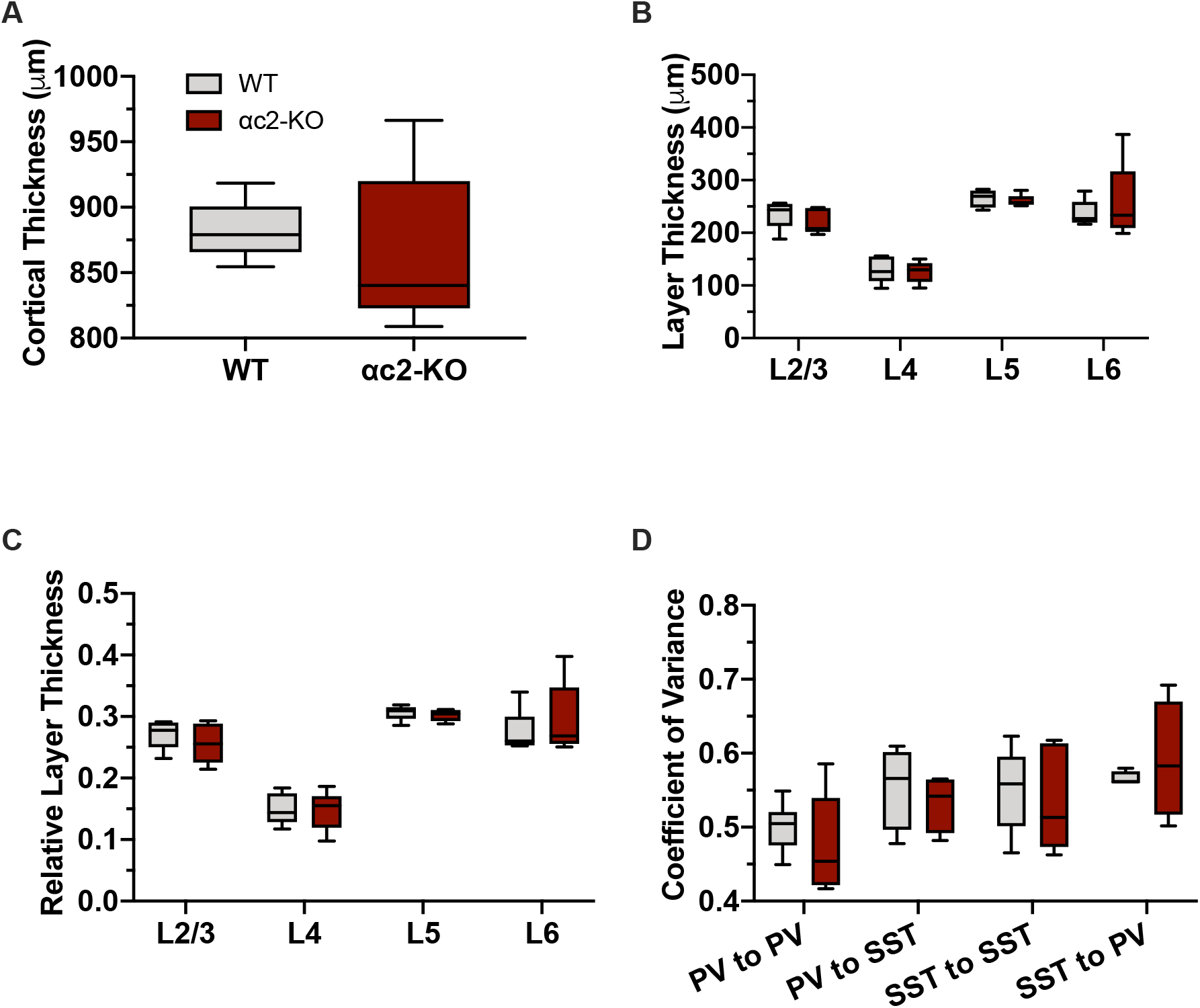
Cortical thickness and CV are unaltered in the αc2-KO. A-Thickness of P30 V1, WT vs. αc2-KO. Cortical thickness is not significantly altered in the αc2-KO (p=0.5832, unpaired T-test with Welch’s correction) B-Laminar thickness of P30 V1, WT vs. αc2-KO. Laminar thickness is not significantly altered in the αc2-KO (LtR: p=0.9342, 0.9989, 0.9992, 0.8140, 2-way ANOVA with SMCT) C-Relative laminar thickness of P30 V1, WT vs. αc2-KO. Proportionally, layers are not significantly altered in the αc2-KO (LtR: p=0.9205, p=0.9998, p=0.9993, p=0.7688, 2-way ANOVA with SMCT) D-CV between PV-PV, PV-SST, SST-SST, SST-PV pairs in P30 V1, WT vs. αc2-KO. CV is not different for any cell pair examined (LtR: p=0.9137, p=0.9949, p=0.9673, p=0.8717, 2-way ANOVA with SMCT)

**Supplemental Figure 10.**
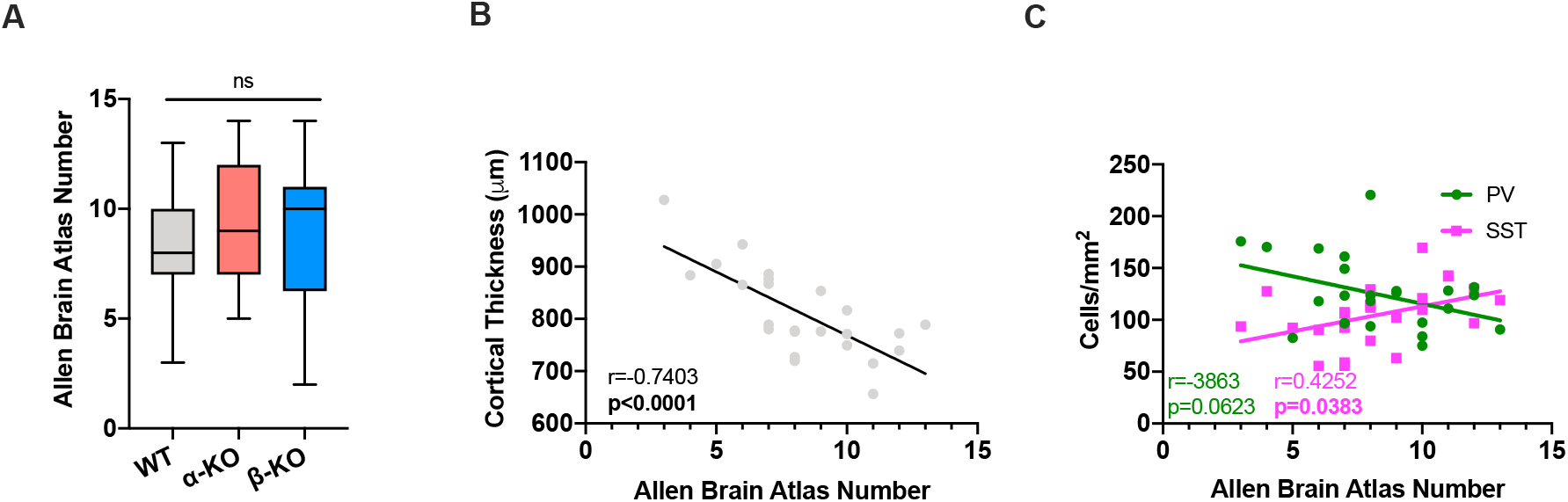
Comparison of Allen Brain Atlas Numbers of images within dataset. A-Comparison of images making up the WT, α-KO, and β-KO datasets. These did not significantly differ in their representation of certain Allen Brain Atlas numbers (ABAN) (LtR: p=0.3243, p=0.5701, p=0.9034, 1-way ANOVA with TMCT) B-Correlation between WT cortical thickness and ABAN. Cortical thickness negatively correlates with ABAN (r=-0.7403, p<0.001, simple linear regression) C-Correlation between WT PV and SST densities and ABAN. PV density is not significantly correlated with ABAN, while SST density is modestly positively correlated with ABAN (PV: r=-0.3863, p=0.0623; SST: r=0.4252, p=0.0383, simple linear regression)

